# Extended role of p21 during the exit from p21-induced G0/G1 arrest

**DOI:** 10.1101/2024.12.23.630045

**Authors:** Dianpeng Zheng, Zhipeng Ai, Suwen Qiu, Yue Song, Chenyang Ma, Weikang Meng, Runju Zhang, Feng He, Hongqing Liang, Jun Ma

## Abstract

Stress- or developmentally-induced signals can trigger G0/G1 cell cycle arrest through the upregulation of p21, a key inhibitor that halts proliferation. While p21 is widely recognized as essential for establishing this arrest, its degradation is generally assumed to be a prerequisite for the exit from arrest and cell cycle re-entry. Using a rapid p21 degradation system that allows for both endogenous p21 tracking and controlled depletion, we uncovered an extended role for p21 during the exit from G0/G1 arrest. Our results showed that removal of the arrest inducing signals did not lead to an immediate decline of p21 during the exit process; instead, p21 levels continued to maintain for a period before its gradual down-regulation. Importantly, premature depletion of p21 during this process weakened the capacity of cell cycle re-entry, particularly in cells with high levels of pre-existent p21. We found that during the release from arrest, cells with high levels of pre-existent p21 but a premature p21 depletion exhibited reduced capacity to restore KRAS/ERK activity, and supplementing KRAS/ERK activity rescued the failure of cell cycle re-entry in these cells. These findings reveal a previously unappreciated function for p21 during the exit from G0/G1 arrest, aside and distinct from its role in initiating the arrest. This new paradigm extends across multiple cell lines and stress-induced arrest contexts, offering a new framework for understanding p21 function in response to anticancer therapies and in development.

## Introduction

In response to a wide range of intra- and extracellular signals such as cellular stress and developmental cues, cells encounter a pivotal decision point of either to proliferate or enter G0/G1 arrest^1, 2^. They enter the G0/G1 arrest state to overcome stress or prepare for reaction to stimuli^3–6^. The primary mechanism triggering G0/G1 arrest is through halting the activation of cyclin D-CDK4/6 complexes and cyclin A/E-CDK2 complexes^7^. The lack of CDK activity restricts cell cycle entry through preventing RB phosphorylation and E2F1 activation^8^. Thus, cells can enter G0/G1 arrest either by a direct loss of pro-proliferative factors in the case of serum starvation, or through up-regulation of cell cycle inhibitors, such as p27 or p21, as a result of various external stress conditions^9, 10^. A widely accepted model posits that cell cycle inhibitors are induced to counteract pro-proliferative signals, acting as a “barrier” to proliferation. It is generally thought that, during the exit from the arrest, the cell cycle network becomes rebounded by the removal of cell cycle inhibitors and a consequent revival of the pro-proliferation signals^11^. Under the framework of this model, cells arrested in G0/G1 are expected, and have been well documented, to revert back to proliferation when the cell cycle inhibitors are removed^12^.

Among cell cycle inhibitors, p21 is the pivotal cyclin-dependent kinase inhibitor mediating the damage/stress-induced or developmental G0/G1 arrest^13–15^. p21 is responsive to various environmental or cellular stresses^16^, having a crucial role in the G0/G1 arrest state of stem cells, including NSCs and HSCs^17, 18^. p21 can be activated by p53, TGF-β/SMAD or PI3K/FOXO pathways^19–21^ to halt cell cycle progression through physically sequestering and inhibiting CDK2 or CDK4/6. The intricate interplay between p21 and CDK activities has been regarded as the key determinant in G0/G1 arrest-proliferation decision^22^. Importantly, genetic knockout studies show that p21 depletion alone is sufficient to abolish stress-induced G0/G1 arrest^15^. Utilizing live-cell p21 reporters to track endogenous p21 levels, it is observed that even low levels of p21 protein can suppress CDK2 activity and effectively induce G0/G1 arrest^13, 14^. Recent studies suggest that p21 levels and its accumulation dynamics exhibit heterogeneity and are tightly associated with cell cycle arrest and recovery fates. For example, upon chemotherapy in cancer cell, it was found that, while cells with intermediate levels of p21 retain the capacity to exit from the arrest upon subsequent drug withdrawal, those that accumulate high or low levels of p21 are driven into senescence^23^. In another study that was based on multiplexed single cell imaging, it was suggested that high p21 levels in combination with pro-proliferative factors can together drive cells into senescence^10^. These studies suggest that p21 is not only important for the arrest maintenance but also bears a potential influence for cell fate determination after the arrest. While traditional studies have been mainly focused on the role of p21 during the arrest, whether and how p21 may contribute to the exit from G0/G1 arrest and the subsequent cell fates remains an open question.

To monitor the dynamics of p21 level and to investigate the role of p21 during the exit from G0/G1 arrest, we constructed a quantitative single-cell live imaging approach in combination with an acute p21 depletion system. Our live cell imaging combined with controlled depletion of p21 during the exit process revealed a previously unappreciated role of p21 in safeguarding the G0/G1 arrest and cell cycle re-entry. Such a role is sensitive to the pre-existent levels of p21. While its premature depletion during arrest release had minimum effect on cells with low p21 levels, it prevented cell cycle re-entry of cells with high p21 levels. We show that such an inability to resume cell cycle for these cells is a result of an impaired reactivation of KRAS/ERK signaling. Thus, our work uncovers a novel role of p21 acting during the exit phase of stress-induced G0/G1 arrest.

## Results

### p21 dynamics in G0/G1 arrest induced by various stress and developmental signals

To simultaneously track the dynamics of p21 and investigate its temporal roles in G0/G1 arrest, we utilized a p21 knock-in system suitable for both live-imaging detection and an induced degradation^24^. We modified both alleles of *CDKN1A*, the p21-encoding gene by endogenously tagging with miniIAA7-mTurquoise2 (Figure S1A-C; see Methods for details), together with the integration of a CDK2 biosensor as indicator for defining G0/G1 arrest-proliferation states^25^, an mMaroon1-tagged H1.0 as nuclear marker, and an mKO2-tagged Cdt1 (aa30-120) as G1/S transition marker^26^ both in hTERT-RPE1 and HL-7702 cells (Figure 1A; S1D, E; also see Methods and legend for details). The cytoplasmic to nuclear fluorescence ratio (Cyt/Nuc) of the mVenus-tagged DHB, which can be directly phosphorylated by CDK2, provides a readout for CDK2 kinase activity^25^.

**Figure 1.**
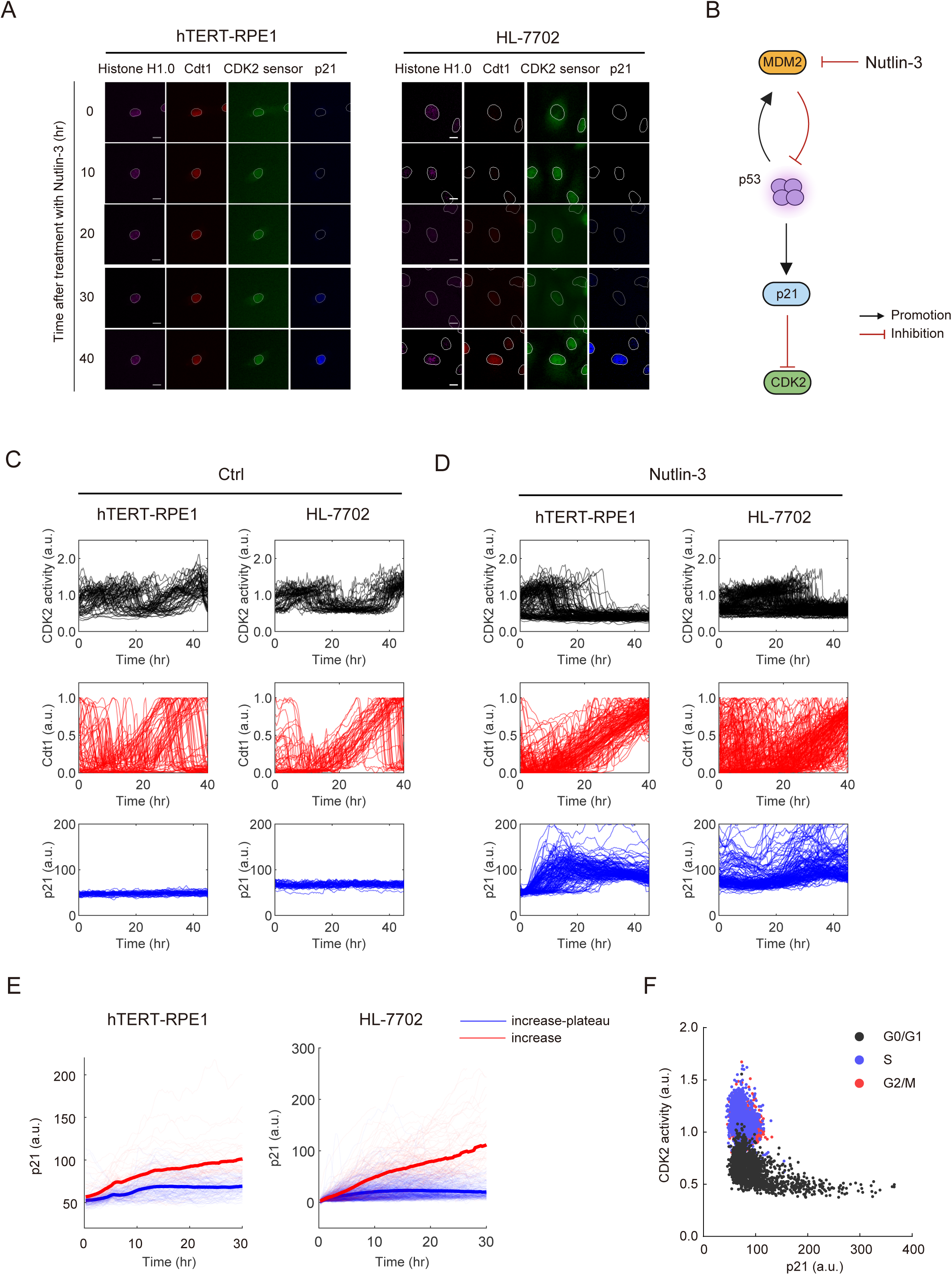
The dynamics of p21 expression induced by various stresses and stimuli. (A) Image showing representative hTERT-RPE1 and HL-7702 p21-AID cells stably expressing the CDK2 sensor, Cdt1, Histone H1.0 and p21-AID, which were treated with 10 μM Nutlin-3 for 0, 10, 20, 30 and 40 hr. Scale bar, 10 μm. (B) Schematics showing p21 activation by Mdm2 inhibitor (Nutlin-3) treatments. (C) Single-cell trajectories showing CDK2 activity, Cdt1 intensity (normalized to maximum) and p21 intensity in hTERT-RPE1 p21-AID and HL-7702 p21-AID cells. (D) Single□ cell trajectories showing CDK2 activity, Cdt1 intensity (normalized to maximum) and p21 levels in hTERT RPE1 p21 AID and HL 7702 p21 AID cells upon treatment with 10□ μM Nutlin□3. (E) Trajectories of single-cell dynamics of p21□ intensity under Nutlin-3 treatment in hTERT□ RPE1 p21□ AID and HL□-7702 p21□-AID cells. The trajectories were aligned to the onset of p21 induction. The population was divided into two groups based on p21 dynamics: red lines indicate p21 trajectories that firstly increasing then followed by plateauing; blue lines show p21 trajectories with continuous increment. The bold line showed the average trajectories for each group. (F) Scatter plot showing the average p21 and CDK2 levels in HL-7702 p21-AID single cells under all treatment conditions. G0/G1, S, G2 phases were marked by black, blue, red colors, respectively.

Several stress-inducing drugs widely used for inducing p21 and G0/G1 arrest were applied (Figure S1F). Specifically, we used Nutlin-3 to directly activate p53 and induce p21 expression (Figure 1B). We also used A-769662 to activate AMPK and mimic nutrient limitation (Figure S1G), topoisomerase II inhibitor Doxorubicin, the DNA double-strand break inducer Zeocin to induce DNA damage, and Palbociclib to induce hypo-osmotic pressure^27^. In addition, we applied TGF-β to induce p21 in an p53-independent manner (Figure S1H) to mimic a transition into p21-meidated G0/G1 arrest mesenchymal state^28–31^.

We performed live cell imaging to track the dynamics of p21, CDK2 activity, Cdt1 level in proliferating cells and upon stress-treatment (Figure 1C, D; S2A, B). In unperturbed condition, the percentage of proliferation exceeded 80% and 70% in mother and daughter cells respectively (Figure S2C, D), with an average cell-cycle length of 28 hr (Figure S2E). Upon the treatment of various stimuli, the cell cycles were significantly elongated, and the proliferation rates were compromised to varying degrees (Figure S2C-E). Across all treatments, we found that p21 accumulation took place only during G0/G1 phase (Figure S2F), and it exhibited a positive correlation with the length of G0/G1 phase (Pearson correlation 0.2512, p<0.0001) (Figure S2G). In contrast, proliferative cells did not exhibit significant accumulation of p21 when passing through their G1 phase despite the heterogeneity in G1 length (Figure S2H, I).

We further analyzed the dynamics of p21 in cells that entered G0/G1 arrest. A majority of cells typically exhibited an increase-plateau accumulation kinetics of p21, while some of the rest cells could continue to accumulate p21 over the course of the observation (Figure 1D, E). We found that both the variations in the external stress levels (Figure S3 and Supplementary Video 1) and the time of accumulation (Figure S2G) contributed to the p21 level heterogeneity during the G0/G1 arrest. A concurrent monitoring of p21 and CDK2 dynamics revealed that p21 accumulation and CDK2 activation exhibited a mutually exclusive relationship (Figure 1F), confirming that a low level of p21 is sufficient to inhibit CDK2 activity as reported previously^14^.

### Quantitative tracking of p21 revealed an elevated and sustained p21 at the onset of arrest exit

To further monitor the dynamics of p21 during the transition from G0/G1 arrest to cell cycle re□entry, we performed live□cell imaging to simultaneously track p21 levels and CDK2 activity following the removal of the arrest inducing signal. We found that upon Nutlin-3 removal, the large majority of cells retained the capacity to reactivate CDK2 within 20 hr after Nutlin-3 removal (hTERT-RPE1 cells: 64.9 ±10.1%; HL-7702: 62.3 ±8.9%) (Figure 2A). Surprisingly, tracking the single cell dynamics of p21 during the arrest release unveiled an unexpected p21 expression pattern upon the drug removal: instead of an expected decline of p21 following the removal of the stimuli, p21 exhibited an increase for about 6 hr (hTERT-RPE1 cells: 7.27 ±3.49% hrs; HL-7702: 9.44 ±5.40 hrs) (Figure 2B, C) at the initial period of the arrest release before its gradual decline and a complete degradation. This initial maintenance of p21 level was observed in both hTERT-RPE1 and HL-7702 cell lines (Figure 2A, B). During the periods of sustained p21 expression, CDK2 activity remained low and stable, suggesting that p21 was actively inhibiting CDK2 (Figure 2A, B). Furthermore, CDK2 activity commenced around the termination of the sustained p21 expression period, rather than upon complete p21 degradation (Figure 2C). This pattern was also observed under different drug treatment conditions (Figure 2D).

**Figure 2.**
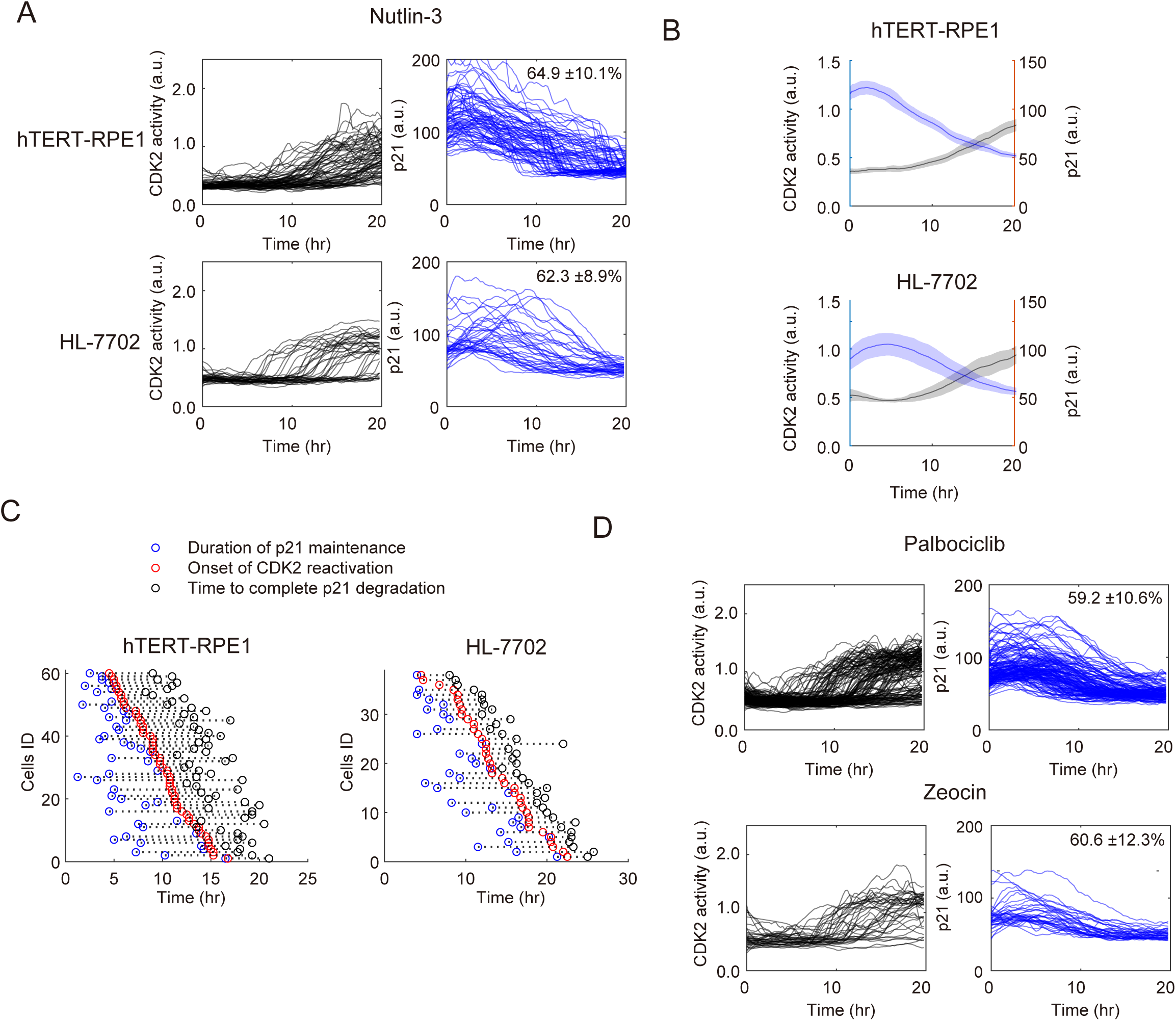
Quantitative tracking of p21 dynamics during the release from G0/G1 arrest. (A) hTERT-RPE1 and HL-7702 cells were treated with Nutlin-3 for 48 hr, followed by release into normal medium. Time-lapse imaging was performed for 20 hr after drug release. Black lines indicate CDK2 activity and blue lines indicate p21 levels. Recovery percentages are noted on the right as mean ± SD. (B) Average trajectories and 95% confidence interval of p21 intensity and CDK2 activity from (A). (C) Scatter plots depicting the initial time of p21 degradation (blue), the onset time of CDK2 reactivation (red), and time of complete p21 degradation (black) at the single□ cell level. Dashed lines connected the values from individual cells. (D) HL-7702 cells were treated with Palbociclib or Zeocin for 48 hr, followed by release into normal medium. Time-lapse imaging was performed for 20 hr after drug release. Black lines indicate CDK2 activity and blue lines indicate p21 levels from individual cells. Recovery percentages are noted on the right as mean ± SD.

Taken together, our results indicate that, while p21 accumulation broadly mediates a reversible G0/G1 arrest under stress, its expression extended to the initial phase of the arrest release, indicating p21 may continue to function during the exit from the G0/G1 arrest.

### Rapid depletion of p21 during arrest release impairs cell cycle re-entry efficiency

We noticed the increase of CDK2 activity only began after p21 level started to decline (Figure 2B, C), therefore we asked whether depleting p21 at the beginning of the exit process would accelerate the speed of CDK2 reactivation. We utilized the p21 knock-in reporter containing the miniIAA7-mTurquoise2 for an induced and rapid degradation of p21 (Figure S4A). In this system, the endogenous p21 protein became undetectable within one hour after the addition of IAA (Figure S4B, C), and this degradation was dependent on an active ubiquitin pathway as expected (see Figure S4C, D and legend for details). At the population level, degradation of p21 reduced the percentages of G0/G1 arrested cells in different G0/G1 arrest-inducing conditions, highlighting the role of p21 in maintaining the G0/G1 arrest (Figure S4E).

We followed single cells over the course of p21 depletion by live imaging. Cells with Nutlin-3 treatment induced and maintained p21 expression in the G0/G1 arrest state (Figure S5A). After 32 hr of Nutlin-3 treatment, we removed Nutlin-3 to induce release from arrest while added in IAA to deplete p21 (Figure 3A; S5B). We found in cells undergoing exit from arrest with p21 depletion, around 30% of cells exited the arrest (Figure 3B, C). This is in sharp contrast with cells that re-entered the cell cycle after drug removal without p21 depletion (hTERT-RPE1 cells: 64.9 ±10.1%; HL-7702: 62.3 ±8.9%) (Figure 2A). At single cells level, distinct CDK2 activity and cell-cycle dynamics can be observed upon p21 depletion during arrest release (Figure 3B, C). Upon p21 depletion, 30% of the cells re-activated their CDK2 and they were referred as the “exit” subgroup (Figure 3B, C). A majority of cells (about 70%) remained a CDK2-inactive state even with a complete depletion of p21 (Figure 3B, C; Supplementary Video 2). We refer to these cells as the “arrest” subgroup. The failure of re-activating CDK2 in a large majority of cells upon p21 depletion suggests a temporal sustenance of p21 after the initiation of the release process is potentially important for the proper resumption of the cell cycle.

**Figure 3.**
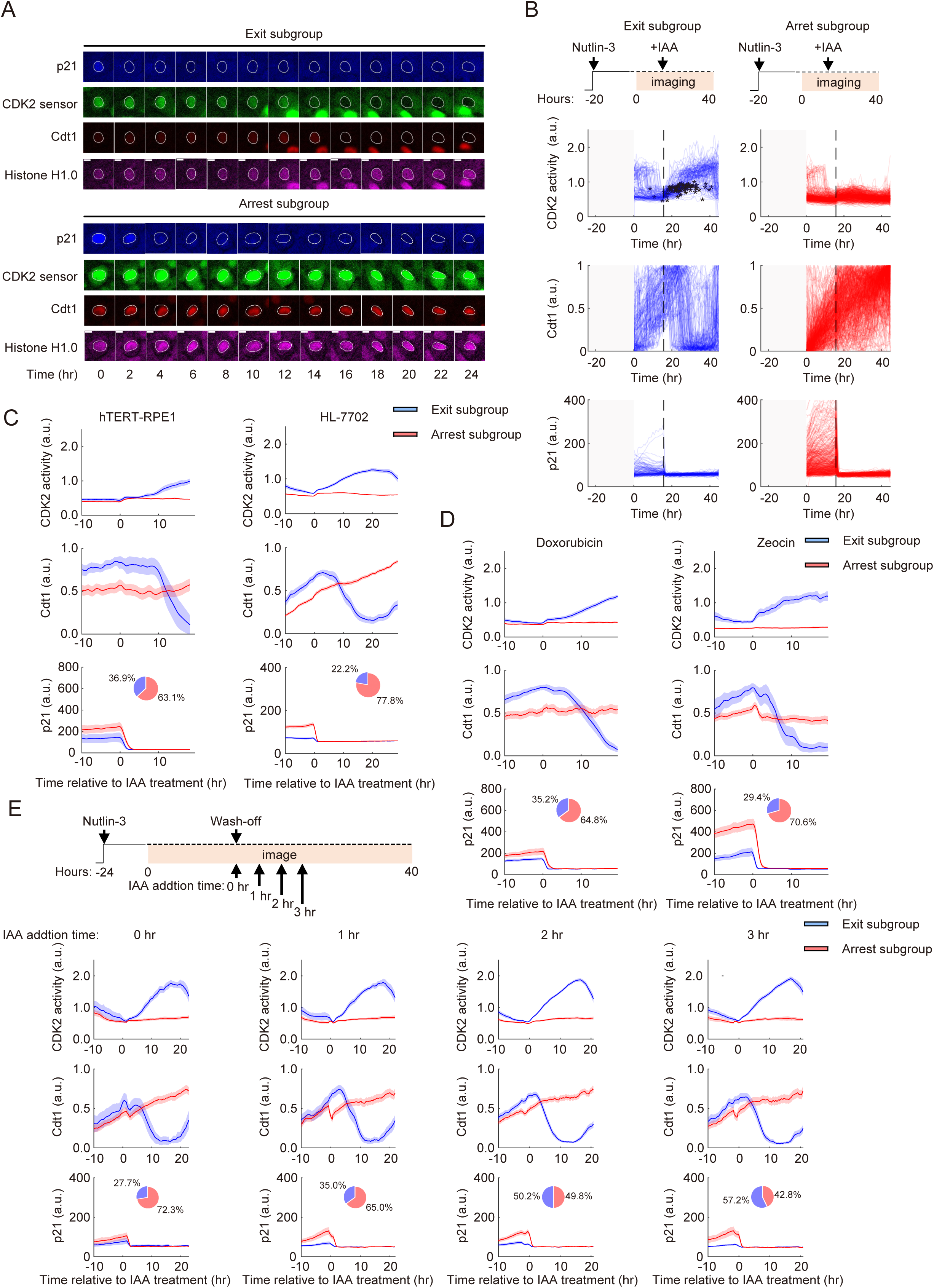
Rapid depletion of p21 during the arrest release compromises efficient cell cycle progression. (A) Single cell fluorescent images showing CDK2 activity, Cdt1 and p21 expression in HL-7702 p21-AID cells treated with 10□ μM Nutlin 3□ for 24□h, followed by IAA addition and Nutlin□3 removal (marked as 0 h). Cells were imaged for 48 hr and divided into two subgroups, “exit” and “arrest”, based on CDK2 trajectories after IAA addition. Scale bar, 10 μm. (B) Single-cell trajectories showing CDK2 activity, Cdt1 intensity (normalized to maximum) and p21 intensity in HL-7702 p21-AID cells. Cells were pretreated with 10□μM Nutlin 3□for 24□h, followed by time□lapse imaging for 12□h prior to the addition of 500□μM IAA and Nutlin□3 removal. Cells were divided into two subgroups, “exit” and “arrest”, based on CDK2 trajectories after IAA addition: blue lines represent cells with increasing CDK2 activity upon p21 removal (“exit”); red lines indicate cells remaining in a CDK2-low state even after p21 removal (“arrest”). Black asterisks indicate G1/S transition. (C) Average trajectories of CDK2 activity and p21 intensity in hTERT-RPE1 and HL-7702 p21-AID cells. Cells were treated with Nutlin-3 (*N*=230; *N*=780) for 24 hr, followed by time□lapse imaging for 12□h prior to Nutlin 3 removal with the addition of 500□μM IAA. Cells were divided into two subgroups, “exit” (blue) and “arrest” (red), based on CDK2 trajectories after IAA addition. Data are presented as mean (solid lines) ± 95% confidence intervals (shaded area). Pie charts illustrate the percentage of cells in different subgroups under the indicated conditions. (D) Average single-cell trajectories of CDK2 activity, Cdt1 intensity (normalized to maximum) and p21 intensity in hTERT-RPE1 p21-AID cells. Cells were treated with Doxorubicin or Zeocin for 24 hr, followed by time□lapse imaging for 12 h prior to the drug removal with addition of 500□μM IAA. Data are presented as mean (solid lines) ± 95% confidence intervals (shaded area). Pie charts illustrate the percentage of cells in different subgroups under Doxorubicin (*N*=397) and Zeocin (*N*=235) treatments. Blue lines represent cells with increasing CDK2 activity upon p21 removal (“exit”); red lines indicate cells remaining in a CDK2-low state even after p21 removal (“arrest”). (E) Average trajectories of CDK2 activity and p21 levels in HL-7702 p21-AID cells. Cells were pre-treated with 10 μM Nutlin-3 for 24 hr, and then subjected to time-lapse imaging for 12 hr before Nutlin-3 removal. At the time of washout, 500□μM IAA was added at 0, 1, 2, or 3□hr, respectively. Cells were divided into two subgroups, “exit” (blue) and “arrest” (red), based on CDK2 trajectories after IAA addition. Data are presented as mean (solid lines) ± 95% confidence intervals (shaded area). Pie charts illustrate the percentage of cells in different subgroups under the indicated conditions.

To evaluate whether the observed failure in cell cycle re-entry upon p21 depletion is a property peculiar to Nutlin-3 induced arrest, we performed live-cell imaging to track the dynamics of G0/G1 arrest induced by a diverse set of stress conditions (Figure S5C). We followed both CDK2 activity and Cdt1 level upon stress removal with p21 depletion and found that, under different treatment conditions, there was always a significant proportion of cells (approximately 70-80%) that failed to reactivate CDK2 after p21 depletion. These cells remained arrested, exhibiting properties of “arrest” subgroups (Figure 3D; S5D). The cell cycle re-entry rate upon release with pre-mature p21 degradation was substantially lower than that after merely removing the arrest inducing signal (Figure 3D; S5D). Together, our results suggest that a role of p21 in safeguarding cell cycle re-entry during the exit from G0/G1 arrest is a general phenomenon, conserved across different cell types and stress conditions.

To define a temporal requirement for p21 during the release process, we depleted p21 at various time points following the removal of arrest inducing signal (Figure 3E). Live-cell imaging revealed that earlier p21 depletion resulted in a lower proportion of cells reactivating CDK2, whereas depletion at progressively later time points corresponded with increasingly higher rates of CDK2 reactivation (Figure 3E). These findings indicate that p21 is critically required during the initial phase of exit from G0/G1 arrest. Its importance diminishes over time (Figure 3E), suggesting that p21 functions primarily in the early window of the release to enable proper cell cycle re-entry potential.

### The requirement for p21 during exit from arrest is associated with the pre-existent p21 level

To determine whether the essential role of p21 during the exit from arrest is related to the progression of the G0/G1 arrest, we initiated release from the arrest by removing the stress-inducing drugs after different periods of treatment. This resulted in the arrested cells accumulating varying levels of p21 prior to release (Figure 4A). When release with p21 depletion was initiated at an early time point of the arrest (25 hr), cells generally exhibited a lower p21 level comparing to that initiated at later time point (36 hr) (Figure 4B; Figure S6A). Regardless of the timing of p21 depletion, the same two fates of CDK2 reactivation dynamics can be observed (Figure 4C; Figure S6B), but their proportions differed dramatically. Early release without p21 exhibited a higher proportion of “exit” cells and fewer “arrest” cells compared to late release with p21 depletion (Figure 4C; Figure S6B). These results are supportive of a stronger dependency of p21 for proper recovery when cells have been in the arrested state for a longer period of time with a higher pre-release p21 level.

**Figure 4.**
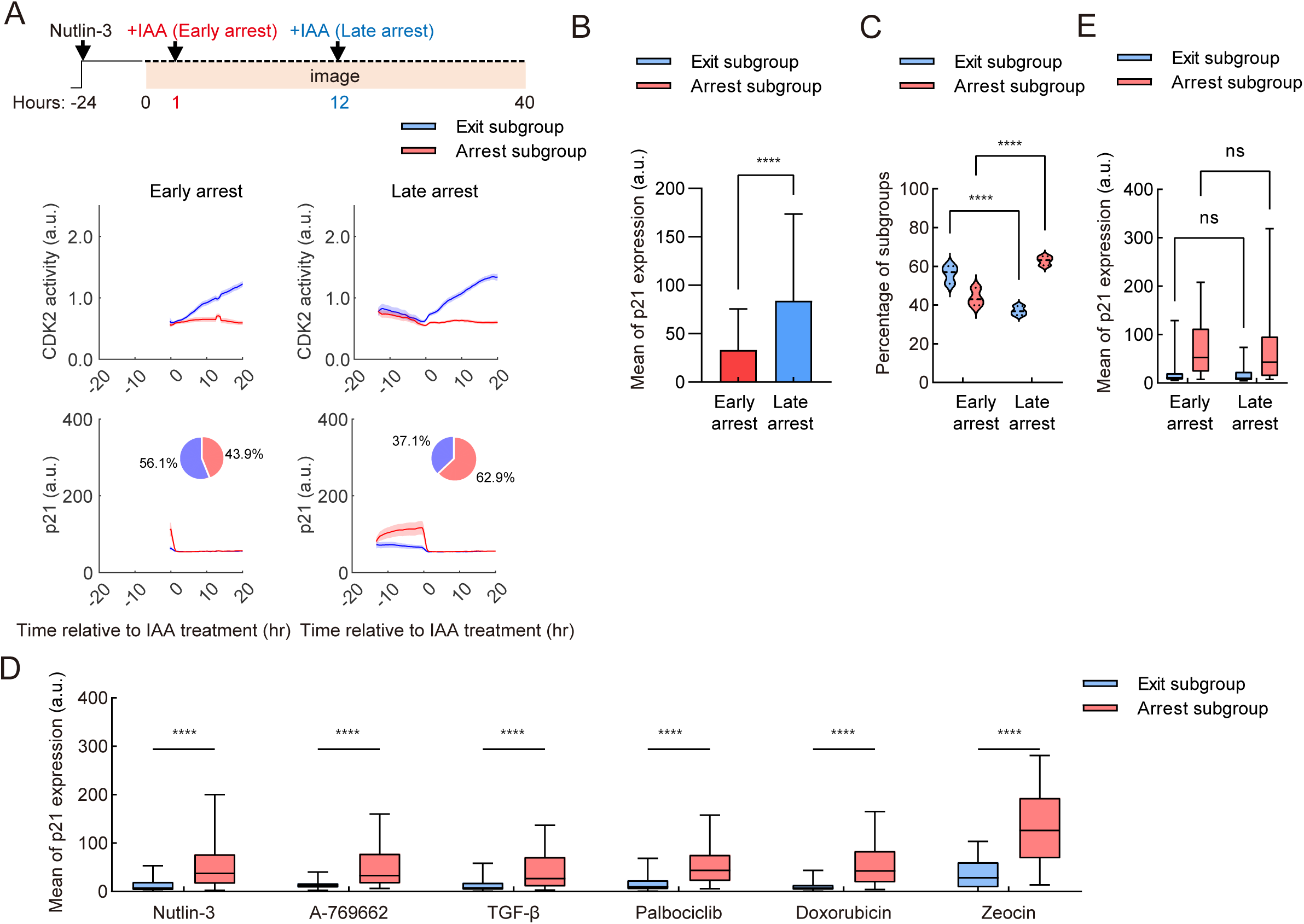
Rapid p21 degradation after different duration of the G0/G1 arrest. (A) Average trajectories of CDK2 activity and p21 levels in HL-7702 p21-AID cells. Cells were pre-treated with 10 μM Nutlin-3 for 24 hr, and then subjected to time-lapse imaging for 1 hr (early arrest, *N*=152) or 12 hr (late arrest, *N*=196) before the addition of 500 μM IAA and Nutlin-3 removal. Cells were divided into two subgroups, “exit” (blue) and “arrest” (red), based on CDK2 trajectories after IAA addition. Data are presented as mean (solid lines) ± 95% confidence intervals (shaded area). Pie charts illustrate the percentage of cells in different subgroups under the indicated conditions. (B) Bar plots illustrating the average p21 levels at different time points before p21 depletion and Nutlin-3 removal (early arrest or late arrest). Two-way ANOVA was performed, “ns” and ** and **** represent p-values > 0.05, < 10^-2^ and < 10^-4^. The error bars represent standard deviation (SD) for the result obtained from three independent live tracking experiments. (C) Violin plots illustrating the percentage of “exit” (blue) and “arrest” (red) subgroups upon p21 depletion and Nutlin-3 removal at early or late time point. Mixed-effects analysis was performed, and “ns”, ** and **** represent p-values > 0.05, < 10^-2^ and < 10^-4^ respectively. (D) Box plots depicting the distribution of average p21 level within 5 hrs’ window before IAA addition and drugs removal in “exit” and “arrest” subgroups under different drug treatment to induced G0/G1 arrest. Mixed-effects analysis was performed and “ns”, *, **, *** and **** represent p-values > 0.05, < 0.05, < 10^-2^, < 10^-3^ and < 10^-4^ respectively. (E) Box plots illustrating the distribution of the mean p21 levels of single cells in the “ exit” (blue) and “arrest” (red) subgroups from the early or late time point of p21 depletion cells during Nutlin-3 treatment. Student’s t-test was performed, and ** and **** represent p-values > 0.05, < 10^-2^ and < 10^-4^ respectively.

Further analysis of single cell heterogeneity supported this notion. We found that the subgroups of single cells with different CDK2 reactivation dynamics were associated with distinct pre-release p21 levels: with the “exit” subgroups exhibiting significantly lower p21 level (<40 a.u.) than the “arrest” subgroup (>80 a.u.) (Figure 4D). Furthermore, with G0/G1 arrest progressing to later time point, we detected an overall higher p21 level and increased proportion of “arrest” cells (Figure 4C; S6B). Importantly, we found that, during release from arrest with p21 depletion, the subgroups of cells with similar p21 levels exhibited remarkably similar fates, regardless of whether release with p21-depletion was induced at early or late arrest time points (Figure 4E; S6C). This relationship was also consistently observed across release from different arrest-inducing drugs (Figure 4D). Thus, for cells with high pre-existent p21 level, p21 is more essential in safeguarding the proper CDK2 reactivation and cell cycle re-entry.

### The temporal maintenance of p21 during the exit from arrest is essential for reviving the cell cycle

We further investigated whether the maintenance of p21 itself, but not other indirect effects, is what safeguards the cell cycle re-entry and CDK2 reactivation following the release from G0/G1 arrest. To address this, we generated hTERT□RPE1 cells with doxycycline□inducible p21□dTAG expression, enabling us to induce G0/G1 arrest though controlled exogeneous p21 expression (Figure 5A). A range of doxycycline concentrations was applied to achieve graded p21 levels, mirroring the expression spectrum observed under different stress conditions (Figure 5B, C). As expected, distinct p21 levels were induced within 24 hr, accompanied by suppression of CDK2 activity, confirming the establishment of p21 mediated G0/G1 arrest (Figure 5B, C). We selected 1 µg/mL and 10 µg/mL doxycycline to model arrest conditions with low and high p21 expression, respectively. After induction, doxycycline was withdrawn to stop p21 expression and allow release from the arrest. Concurrently, p21 was degraded by dTAG addition upon release form the arrest. The cell cycle re-entry rate was then compared between cells subjected to doxycycline washout alone and those with washout and dTAG mediated p21 degradation. While most cells re entered the cell cycle after doxycycline removal, rapid p21 degradation under the high p21 condition severely impaired CDK2 reactivation (Figure 5D, E). In contrast, under low-p21 arrest induced by 1ug/mL doxycycline, p21 depletion during release from arrest did not prevent majority of cells from reactivating CDK2 (Figure 5D, E). These results further confirmed p21 is the essential factor safeguarding the cell cycle re-entry after arrest, and that the pre-existent p21 level prior to arrest release dictates the extent to which p21 is required for the proper exit from G0/G1 arrest.

**Figure 5.**
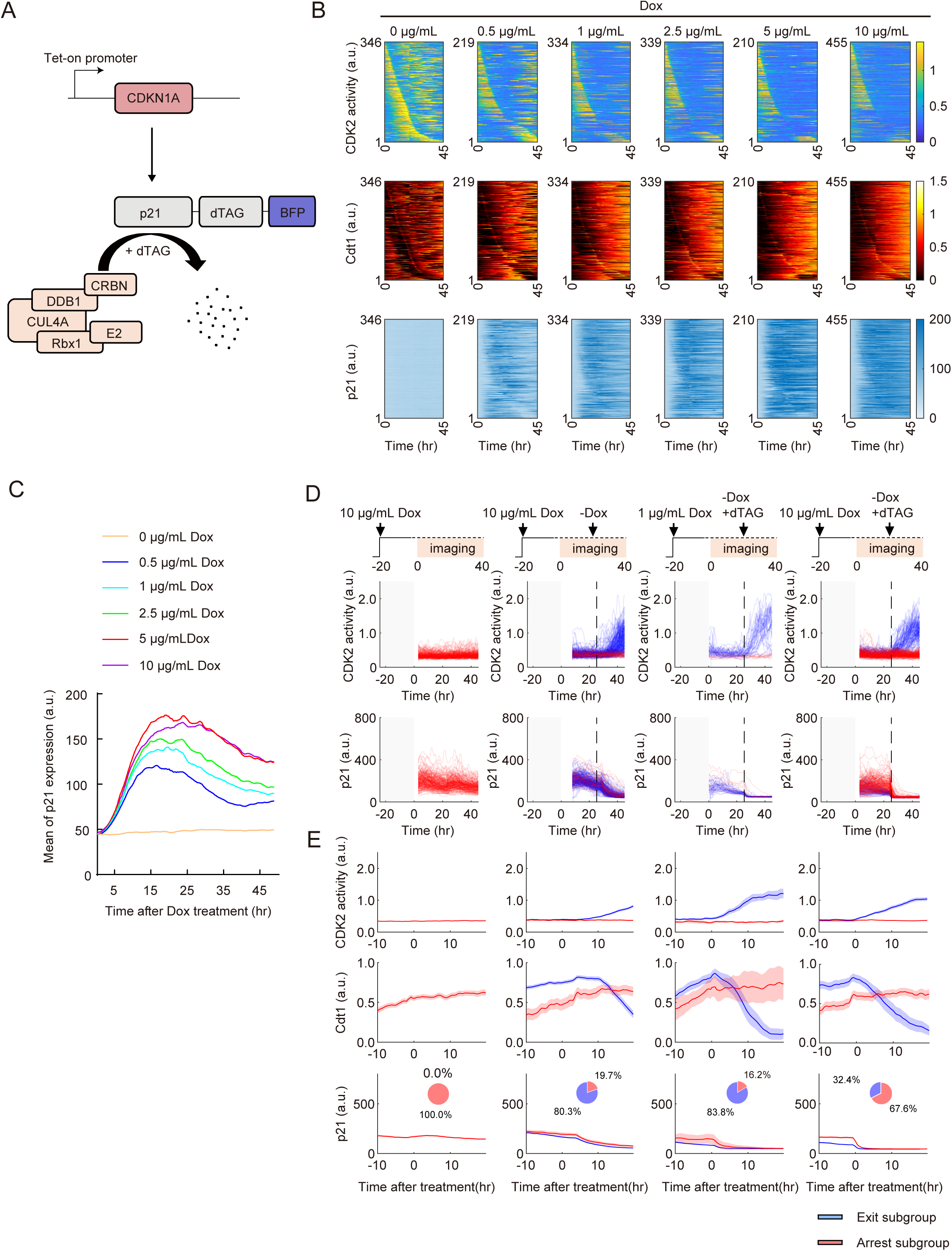
The temporary sustenance of p21 upon release from arrest is critical for reactivating the cell cycle. (A) Schematic of the dTAG-inducible p21 degradation system engineered in hTERT□RPE1 p21-AID cells. Exogenous p21 overexpression is induced by doxycycline. Upon addition of dTAG, the E3 ligase CRBN recognizes and ubiquitinates the dTAG tag fused to p21, resulting in targeted proteasomal degradation. (B) Heatmaps showing single-cell dynamics of CDK2 activity, Cdt1 intensity (normalized to maximum), and p21 levels in response to varying doses of doxycycline combined with IAA over 45 hr. Cells were aligned according to the timing of their first mitotic division. (C) Line plots showing the distribution of mean p21 levels under increasing doses of doxycycline, co□treated with IAA for 45 hr. (D, E) Single-cell trajectories (D) and average trajectories (E) showing CDK2 activity and p21 levels during live-cell imaging of hTERT-RPE1 cells overexpressing p21-dTAG. Cells were pretreated with 500 μM IAA and 10 μg/mL doxycycline for 48 hr, followed by time-lapse imaging for 12 hr prior to doxycycline removal. For the dTAG treatment groups, cells were pretreated with 500 μM IAA and 1 or 10 μg/mL doxycycline for 24 hr, imaged for 12 hr, and then subjected to 500 μM IAA and 500 μM dTAG upon doxycycline removal. Vertical black arrows and dotted lines indicate the timing of different treatments. Cells were divided into two subgroups based on CDK2 activity after p21 depletion, with each subgroup shown in a distinct color. In (E), solid lines represent the mean and shaded areas indicate the 95% confidence intervals. Pie charts show the percentage of cells in each subgroup.

### p21-high cells exhibit reduced CDK4/6 re-activation and ERK pathway gene expression

To identify the mechanism underlying the lost cell cycle re-entry potential due to p21 pre-mature depletion during the exit from the G0/G1 arrest, we firstly determined whether other CDK inhibitors, such as p27, prevented the exit upon p21 depletion. We found the protein expression level of p27 was low in p21-induced G0/G1 arrest state (Figure S6D, E). Furthermore, RNA expression of CDKN1B (p27) also did not exhibit any increment upon p21 induction (Figure S6F), ruling out the possibility of p27 in sustaining the arrest upon p21 removal.

To identify molecular features associated with the inability to reactivate CDK2 upon p21 depletion in p21-high cells, we sorted cells (induced by Palbociclib or Nutlin-3) into p21-negative, p21-low, and p21-high populations (Figure 6A). p21 is a known inhibitor of multiple CDKs, including CDK4/6^32^, CDK2^25^, and CDK1^33^. Co-immunoprecipitation showed that while p21 bound CDK2 in both p21-low and -high groups, its interaction with CDK4/6 was exclusive to the p21-high group (Figure 6B, C). Functional studies tracking CDK4/6 activity sensors in live imaging^32^ confirmed that only p21-high cells showed a pronounced decrease in CDK4/6 activity (Figure S7A-C). However, during release from arrest without p21, the “arrest” subgroup failed to reactivate CDK4/6 (Figure S7D); consequently, this led to a further decrease of pRB in p21-high cells during exit from arrest upon p21 depletion (Figure 6D). The failure to reactivate CDK4/6 during release without p21 points to a fundamental breakdown of the pro-proliferative signaling network necessary for cell cycle re-entry after rapid p21 degradation.

**Figure 6.**
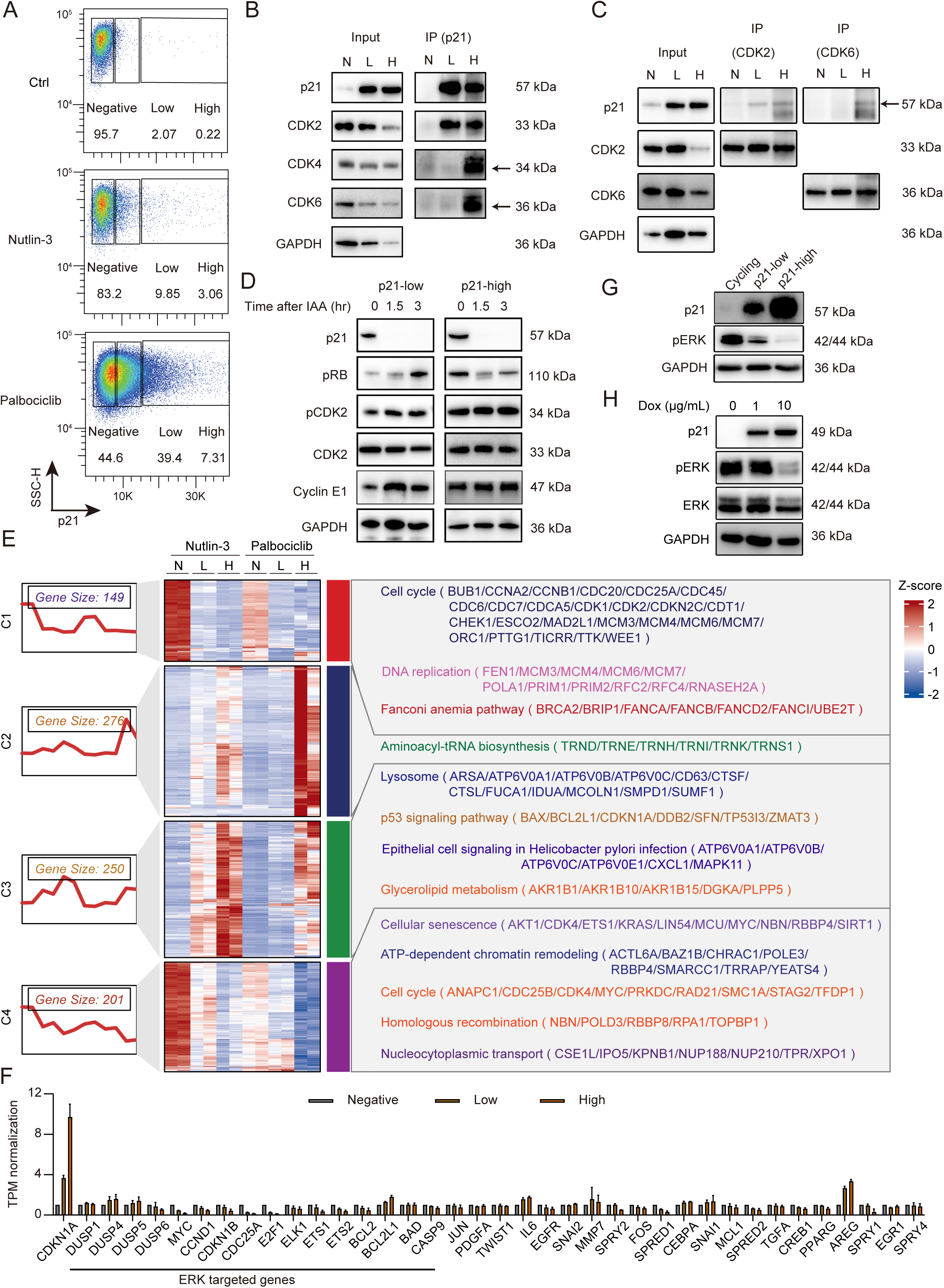
High p21 levels impair the pro-proliferation signals during G0/G1 arrest. (A) Scatter plot showing the endogenous p21 levels (X-axis) and FSC-H (Y-axis) as analyzed by flow cytometry after 48 hr of DMSO, 10 μM Nutlin-3 or 5 μM Palbociclib treatment. The gates were set to separate cells with no p21 expression (negative), low p21 expression (twice the expression of the N group), high p21 expression. The percentages of cells under every gate were noted on the graph. (B, C) Co-immunoprecipitation to detect the interaction between p21, CDK2, CDK4, and CDK6 proteins in HL-7702 p21-AID cells after 48 hr of 10 μM Nutlin-3 treatment. Cells with negative p21 (N), low p21 (L) and high p21 (H) expression were sorted out from FACS as shown in (A). CDK2, CDK4, or CDK6 was pulled down to detect the co-precipitation of p21 (B). Conversely, p21 was pulled down to detect the co-precipitation of CDK2 or CDK6 (C). (D) Western blot images showing the p21 level, RB phosphorylation at Ser807/811, CDK2 phosphorylation at Thr160, CDK2 and Cyclin E1 levels in cells with different p21 expression level before its depletion. The cells treated with 10 μM Nutlin-3 for 48 hr were sorted into p21-low (Low) and p21-high (High) groups under the same gating as shown Figure 2A. After washout, the “low” and “high” cells were treated with 500 μM IAA to deplete p21 and samples were collected at 0, 1.5, and 3 hr for protein expression analysis. (E) Integrated heat map (log_2_ fold change), KEGG enriched pathways and representative differentially expressed genes in control (N), p21-low (L) and p21-high (H) cells under 10 μM Nutlin-3 or 5 μM Palbociclib treatment. Genes were grouped into four clusters based on expression dynamics under different conditions. (F) Bar plots showing TPM-normalized expression levels of ERK related genes in control (N), p21-low (L), and p21-high (H) cells under 10 μM Nutlin-3 treatment. (G) Western blot images showing the expression of p21, phospho-ERK and GAPDH in HL-7702 p21-AID cells treated with 10 μM Nutlin-3 for 48 hr. p21-low and p21-high cells were sorted by flow cytometry based on gating in (A). (H) Western blot images showing the expression of p21, phospho-ERK, ERK and GAPDH in hTERT-RPE1 p21 overexpression cells treated with 500 μM IAA and 0, 1 and 10 μg/mL doxycycline for 48 hr, respectively.

To elucidate the mechanism behind the CDK4/6 reactivation failure upon p21 depletion in p21-high cells, we performed RNA-seq sorted by p21 levels as well (Figure 6E). We identified genes whose expression correlated with p21 levels. Notably, p53 pathway and lysosome pathway genes showed a positive correlation with p21^34^. Conversely, genes negatively correlated with p21 (cluster 4 genes) were enriched for cell cycle and senescence pathways (Figure 6E). Importantly, key pro-proliferative driver genes like KRAS and MYC, critical for ERK signaling and CDK4/6 activation, were significantly downregulated in p21-high cells. This was further evidenced by a broad downregulation of ERK target genes specifically in this population (Figure 6F). We also checked the phospho-ERK level and found that phospho-ERK was dramatically down-regulated in cells with high p21 expression (Figure 6G, H), consistent with the RNA-seq data. This distinct transcriptomic profile in p21-high cells indicates a systematic weakening of the mitogenic signaling network in these cells. This may pose additional barrier for re-activating CDK4/6 and promoting cell cycle re-entry during the exit from the arrest.

### ERK activation failure underlies the cell cycle re-entry defects during arrest release without p21

To evaluate whether pre-mature p21 depletion affected the ERK signalling during the exit from arrest, we further checked the phospho-ERK level during the release upon p21 depletion in p21 high and low sub-populations. We found there was an exacerbated loss of pERK upon p21 depletion in p21 high cells (Figure 7A). These results implied that a temporal sustenance of p21 favoured the maintenance of ERK activity during exit from the G0/G1 arrest. To further confirm this, we checked the phospho-ERK level during the release from p21-overexpression induced G0/G1 arrest.

**Figure 7.**
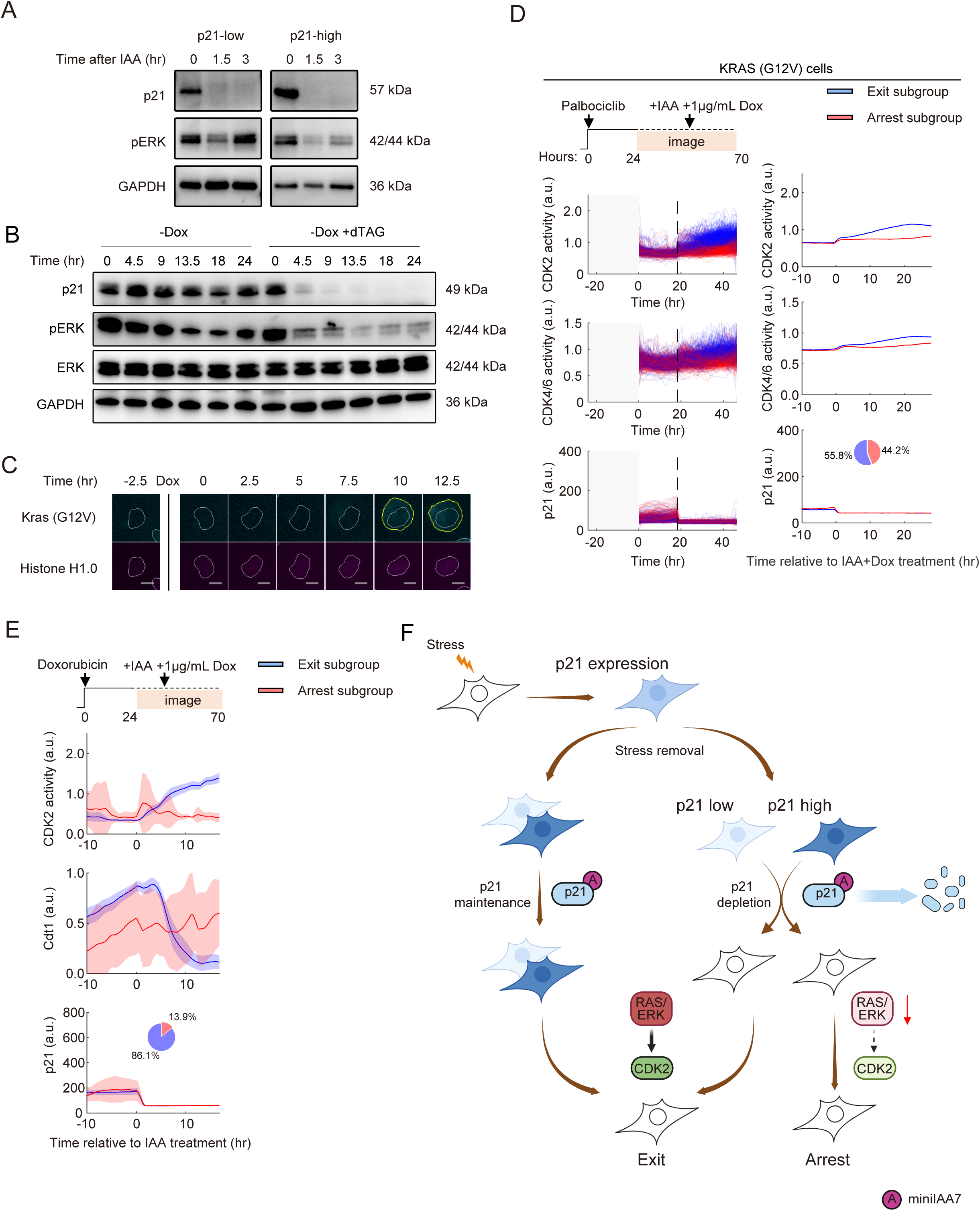
Failure to maintain ERK activity underlies the cell cycle re entry defects during arrest release without p21. (A) Western blot images showing p21 level, phospho-ERK and GAPDH in HL-7702 p21-AID cells treated with 10 μM Nutlin-3 for 48 hr. p21-low and p21-high cells were sorted by flow cytometry based on gating in Figure 6A. After washout, sorting cells were treated with 500 μM IAA and protein samples were collected at 0, 1.5 and 3 hr, respectively. (B) Western blot images showing p21 level, phospho-ERK, ERK and GAPDH in hTERT-RPE1 p21 overexpression cells treated with 500 μM IAA and 10μg/mL doxycycline for 48 hr. Subsequently treated without or with 500 μM dTAG upon doxycycline removal and protein samples were collected at 0, 4.5, 9, 13.5, 18 and 24 hr, respectively. (C) Representative fluorescent images showing the induction of BFP-KRAS (G12V) upon addition of 1 μg/ml doxycycline at 0 hr in HL-7702 p21-AID cells stably expressing Tet-on KRAS (G12V). Scale bar, 20 μm. (D) Single-cell trajectories (left) or average trajectories (right) showing CDK2, CDK4/6 activity and p21 level during live cell imaging of HL-7702 p21-AID cells with KRAS (G12V) overexpression (*N*=208) upon p21 depletion. Cells were pre-treated with 5 μM Palbociclib for 24 hr, and then subjected to time-lapse imaging for 12 hr before the addition of 500 μM IAA and 1 μg/ml doxycycline, together with Palbociclib removal. Vertical black arrows and dotted line indicate the time of IAA and doxycycline addition. Cells were divided into two subgroups based on CDK2 activity after p21 depletion with every subgroup shown in different colors. Right panels represented mean (solid lines) ± 95% confidence intervals (shaded area). Pie plots show the percentage of the two subgroups. (E) Average single-cell trajectories of CDK2 activity and p21 levels during live-cell imaging of hTERT-RPE1 p21-AID cells with KRAS (G12V) overexpression upon p21 depletion. Cells were pre-treated with 50 nM doxorubicin (A) (*N*=95) for 24 hr, followed by time-lapse imaging for 12 hr prior to the addition 500 μM IAA and 1 μg/mL doxycycline, together with doxorubicin removal. Vertical black arrows and dotted lines indicate the timing of drug additions. Cells were divided into two subgroups based on CDK2 activity following p21 depletion, with each subgroup shown in different colors. Pie charts depict the proportions of cells in the two subgroups. (F) A schematic diagram illustrating the extended role of p21 during the exit from p21-induced G0/G1 arrest.

Similarly, depletion of p21 during doxycycline withdrawal led to a more dramatic reduction of phospho-ERK (Figure 7B). Thus, the temporal sustenance of p21 during the exit from arrest is important to facilitate the maintenance of ERK activity after the arrest.

To evaluate whether exogenous RAS/ERK reactivation can rescue the inability to re-activate CDK2 after the arrest upon p21 depletion, we engineered a doxycycline-inducible system expressing a constitutively active KRAS□(G12V) mutant, enabling controlled reactivation of this pathway^35, 36^. The G12V mutant provides high KRAS activity independent of external growth factor sensing and transduction. Time-lapse imaging and western blot confirmed robust KRAS□G12V induction within 8□hr of doxycycline addition (Figure 7C; Figure S8A). In G0/G1 arrest cells, KRAS (G12V) expression upon p21 depletion reduced the percentage of the “arrest” subgroups, while increasing the proportion of the “exit” subgroup to more than 50% (Figure 7D, E). This effect was dose-dependent, with CDK2 recovery rates reaching about 65% at higher doxycycline concentrations (Figure S8B, C). The ability of KRAS (G12V) to rescue the cell cycle re-entry upon p21 depletion was also observed under Nutlin-3 induced G0/G1 arrest (Figure S8D-F). These results further suggest the attenuation of RAS/ERK pathway could be the potential mechanism underlying the cell cycle re-entry upon rapid p21 depletion.

Elevating serum levels has been conventionally used to increase the mitogenic signal^37^. We thus checked whether increasing serum concentration can help to restore the CDK2 recovery potential (Figure S9A). Firstly, an increase in serum concentration from 0% to 20% indeed elevated the level of phospho-CDK2, phospho-RB, and cyclin D1 over time (Figure S9B). Then, during release from G0/G1 arrest induced by Nutlin-3, we increased serum concentration from 10% to 20% upon depleting p21 with IAA. This led to an increased expression of phospho-CDK2, phospho-RB, and cyclin D1, as well as mitotic marker cyclin B1 (Figure S9C). Next, we tracked the CDK2 activity in live at single cell level upon supplying 20% of serum at different time points during arrest release with p21 depletion (Figure S9D-F). CDK2 activity in “arrest” subgroups was observed to increase upon elevating serum at different time points (Figure S9D-F). Furthermore, this effect was dependent on ERK activity, as the inhibition of ERK by Ulixertinib reduced the effect of serum elevation (Figure S9G).

Collectively, these results suggested that in p21 is important in safeguarding the RAS/ERK pathway activity to ensure the cell cycle re-entry potential.

## Discussion

G0/G1 arrest is a temporary exit from the proliferative cell cycle. This arrest is generally thought to be governed by either a direct loss of pro-proliferative factors in the case of serum starvation, or through up-regulation of cell cycle inhibitors, p21, in the presence of stresses, with the latter case being more pertinent to G0/G1 observed under physiological or pathological conditions^22^. It has been reported that stress-induced G0/G1 arrest is critically dependent on p21^22^. Even low levels of p21 are sufficient to induce G0/G1 arrest by selectively binding to CDK2 and competing with cyclin A/E^13^. Several recent studies point toward the existence of differential G0/G1 arrest states in tumor^38^ or stem cells^39, 40^. In contrast to its canonical function as a cell cycle inhibitor during G0/G1 arrest, a role of p21 in facilitating or regulating cell cycle re-entry remains largely unknown. An important finding of our analysis is that, after the removal of the stresses, p21 levels remained sustained for several hours before undergoing gradual degradation both in hTERT-RPE1 and HL-7702 cell lines. While many studies have reported that p21 is typically degraded via the ubiquitin ligases CRL4^Cdt2^ and SCF^Skp2^ during G1/S transition^15^, its persistence during early release from the G1/G0 arrest raised the possibility that the sustained p21 expression could be actively function during the release process. Using a precision p21□degradation system in combination with multiple cell□cycle sensors, our live□cell imaging data demonstrate that the sustained presence of p21 during arrest release functions to facilitate the restoration of cell cycle potential.

The role of p21 during stress-induced G0/G1 arrest is through inhibition of proliferation machineries, and consequently, its removal is traditionally expected to lead to cell cycle reactivation^41^. Our work reveals that this textbook model may not be true during the exit from the arrest state. Using an AID system that enables rapid degradation of endogenous p21, we found that in cells with high p21 (“arrest” subgroup), p21 depletion failed to trigger cell cycle re-entry, and this effect was linked to attenuated ERK signaling. Further analysis showed that sustenance of KRAS/ERK signaling during cell cycle re□entry actually requires a temporal p21 expression at the early phase of arrest exit; while p21 became less important at later time point after the releasing from arrest state. Together, these results define a context□dependent role for high p21 in safeguarding cell cycle re-entry (Figure 7F).

KRAS, a proto-oncogene and small GTPase, plays a crucial role in regulating signaling pathways that determine the G0/G1 arrest-proliferation decision^35^. External mitogens activate the RAS/ERK pathway, promoting cell cycle entry. Elevated KRAS activity can drive G0/G1 arrest cells into S phase, while KRAS depletion prevents this transition^42^. Interestingly, CDK2 inhibition in anti-tumor therapies often results in only a transient suppression of CDK2 activity, followed by a rebound within several hours due to residual CDK4/6 activity sustained by the RAS/ERK pathway^43^. p21 is well established as a CDK2 inhibitor, and its expression induces G0/G1 arrest. Accordingly, rapid p21 degradation upon exit from arrest would be expected to facilitate cell cycle rebound via RAS/ERK signaling. However, our data reveal that premature p21 depletion during exit from arrest impairs the restoration of KRAS/ERK signaling. As previous studies have shown, even in proliferating cells, proper cell cycle rebound requires a temporal window for signaling reconstruction^43^. Thus, we propose that full restoration of KRAS/ERK signaling depends on temporal p21 expression during the exit from arrest. This previously unrecognized role of p21 during the exit from arrest offers a new perspective for anti-cancer therapy.

In summary, our study reveals an extended role for p21 in cell cycle re-entry, a role that is distinct from its role in initiating arrest. To address this previously unexplored function of p21, we established an integrated method combining quantitative live-cell imaging with an acute AID-mediated degradation system. This approach allowed us to delineate how p21 differentially controls arrest maintenance and exit. These insights not only refine the mechanistic understanding of stress-induced G0/G1 arrest but may also inform novel therapeutic strategies aimed at modulating cell cycle re-entry.

## Acknowledgment

This study was supported by the National Key R&D Program of China (2023YFC2705601, 2021YFC2700403 and 2018YFA0800102), National Natural Science Foundation of China (32470783, 31871249 and 31871452), and Zhejiang University Basic Research Funding, 226-2022-00206 (H.L.). We acknowledge support of Zhejiang University, Zhejiang University School of Medicine affiliated Women’s Hospital and Children’s Hospital, Fourth Affiliated Hospital of School of Medicine, and International School of Medicine, International Institutes of Medicine, Zhejiang University. We thank Jiajia Wang and Xin Shen from the Core Facilities, Zhejiang University School of Medicine for their technical support. DHB-mVenus was a gift from Tobias Meyer & Sabrina Spencer (Addgene plasmid # 136461; http://n2t.net/addgene:136461; RRID: Addgene_136461). pSH-EFIRES-P-*At*AFB2 was a gift from Elina Ikonen (Addgene plasmid # 129715; http://n2t.net/addgene:129715; RRID: Addgene_129715). pSH-EFIRES-B-Seipin-miniIAA7-mEGFP was a gift from Elina Ikonen (Addgene plasmid # 129719; http://n2t.net/addgene:129719; RRID: Addgene_129719).

## Author contributions

D.Z., F.H., H.L., and J.M. conceived the study and designed the experiments; D.Z., S.Q., Y.S., Z.A., C.M., W.M. and Z.R performed experiments and generated data; D.Z. analyzed the data and generated all figures; R.Z., H.L. and J.M. acquired funding; D.Z., F.H., H.L., and J.M. wrote the paper and all approved the paper.

## Materials availability

Further information and requests for resources and reagents should be directed to and will be fulfilled by the lead contact, Hongqing Liang.

## Data and code availability

Data: The authors declare that all data supporting the findings of this study are available within the paper and its supplementary information. RNA sequence raw data is deposited at SRA (http://www.ncbi.nlm.nih.gov/sra) with the BioProject ID: PRJNA1174361. Data are also available upon request.

Code: RNA-seq analyses were performed using standard software packages. Code for live imaging data analysis is available upon request.

## Experimental model and subject details

### Cell culture

HEK293T and HL-7702 cells were maintained in DMEM (11965092; Thermofisher) supplemented with 10% FBS (SE100-B; VisTech), penicillin/streptomycin (100 U/ml each) at 5% CO2 and 37°C. Media were changed every 2 days and passaged every 2–3 days. To induce epithelial-mesenchymal transition (EMT), HL-0077 cells were treated with 10 μg/ml TGF-β, with media changes every other day. hTERT-RPE1 cells were maintained in DMEM/F12 (1132003; Thermofisher) supplemented with 10% FBS, penicillin/streptomycin (100 U/ml each) at 5% CO2 and 37°C. Media were changed every 2 days and passaged every 2–3 days.

## Method details

### Generating p21-miniIAA7-mTurquoise2 hTERT-RPE1 and HL-7702 cell lines

The generation of endogenously tagged p21-miniIAA7-mTurquoise2 cells was as reported in our previous studies^24^. In brief, the endogenous *CDKN1A* gene was tagged at the C terminus using CRISPR-mediated gene tagging. The gRNA for *CDKN1A* nearby the stop codon was used. The p21-gRNA plasmid (5’-GGAAGCCCTAATCCGCCCAC-3’) and the linearized PCR product of p21-miniIAA7-mTurquoise2-IRES-pruoR donor sequence was transfected into HL-7702 cells at the ratio of 1:1 using jetPRIME, according to the manufacturer’s instructions (101000046; Polyplus). Cells were treated with 10 μg/ml puromycin until outgrown of the colonies. Finally, homozygous knock-in clones were verified by PCR and western blotting.

### Generating *At*AFB2 hTERT-RPE1 and HL-7702 (p21-AID) stable cell lines

HEK293T cells at approximately 70% confluence were transfected with psPAX2, pMD2.G, FUGW-*At*AFB2-mCherry-weak NLS at ratio 0.75:0.5:1 using jetPRIME (101000046; Polyplus), according to the manufacturer’s instructions. Two days after transfection, supernatant was filtered through a 0.45 μm polyethersulfone filter and was used to infect p21-miniIAA7-mTurquoise2 hTERT-RPE1 and HL-7702 cells. After 7 days, mCherry-positive cells were sorted out via flow cytometry (Moflo Astrios EQ; Beckman).

### Generating DHB-Cdt1(30-120)-H1 hTERT-RPE1 and HL-7702 p21-AID stable cell lines

HEK293T cells at approximately 70% confluence were transfected with psPAX2, pMD2.G, and FUGW-DHB-mVenus or FUGW-H1-mMaroon1-IRES-mKO2-Cdt1(30-120) at the ratio of 0.75:0.5:1 using jetPRIME (101000046; Polyplus), according to the manufacturer’s instructions. Two days after transfection, supernatant was filtered through a 0.45 μm polyethersulfone filter and was used to infect hTERT-RPE1 and HL-7702 p21-AID cells. After 7 days, mVenus- and mMaroon1-positive cells were sorted out via flow cytometry.

### Generating 53BP1 DHB-Cdt1(30-120)-H1 HL-7702 p21-AID stable cell lines

HEK293T cells at approximately 70% confluence were transfected with psPAX2, pMD2.G, and FUGW-BFP-truncated 53BP1 (1220-1771) at ratio 0.75:0.5:1 using jetPRIME (101000046; Polyplus), according to the manufacturer’s instructions. Two days after transfection, supernatant was filtered through a 0.45 μm polyethersulfone filter and was used to infect DHB-Cdt1(30-120)-H1 HL-7702 p21-AID cells. After 7 days, BFP-positive cells were sorted out via flow cytometry.

### Generating CDK2 and CDK4/6 sensors HL-7702 p21-AID stable cell lines

HEK293T cells at approximately 70% confluence were transfected with psPAX2, pMD2.G, and FUGW-DHB-mVenus or FUGW-H1-mMaroon1-IRES-mKO2-RB at the ratio of 0.75:0.5:1 using jetPRIME (101000046; Polyplus), according to the manufacturer’s instructions. Two days after transfection, supernatant was filtered through a 0.45 μm polyethersulfone filter and was used to infect HL-7702 p21-AID cells. After 7 days, mVenus- and mMaroon1-positive cells were sorted out via flow cytometry.

### Generating KRAS hTERT-RPE1 and HL-7702 p21-AID stable cell lines

HEK293T cells at approximately 70% confluence were transfected with psPAX2, pMD2.G, and PCW 57.1-BFP-KRAS(G12V) at ratio 0.75:0.5:1 using jetPRIME (101000046; Polyplus), according to the manufacturer’s instructions. Two days after transfection, supernatant was filtered through a 0.45 μm polyethersulfone filter and was used to infect DHB-Cdt1(30-120)-H1 hTERT-RPE1 and HL-7702 p21-AID or CDK2 and CDK4/6 sensors HL-7702 p21-AID cells. After 7 days, cells were treated with 1 μg/ml Doxycycline for 24 hr and then BFP-positive cells were sorted out via flow cytometry.

### Generating p21-dTAG hTERT-RPE1 p21-AID stable cell lines

HEK293T cells at approximately 70% confluence were transfected with psPAX2, pMD2.G, and PCW 57.1-p21-dTAG-BFP at ratio 0.75:0.5:1 using jetPRIME (101000046; Polyplus), according to the manufacturer’s instructions. Two days after transfection, supernatant was filtered through a 0.45 μm polyethersulfone filter and was used to infect DHB-Cdt1(30-120)-H1 hTERT-RPE1 p21-AID cells. After 7 days, cells were treated with 1 μg/ml Doxycycline for 24 hr and then BFP-positive cells were sorted out via flow cytometry.

### Chemical regents

Drugs used in this study include: TGF-β (HY-P7118; MedChemExpress), A-769662 (S2697; Selleck), Palbociclib (S4482; Selleck), Nutlin-3 (S1061; Selleck), Doxorubicin (S1208; Selleck), Zeocin (R25001; Thermofisher), auxin (I5148; Sigma), Doxycycline (ST039A; Beyotime), dTAG (SML2601; Sigma) and Ulixertinib (S7854; Selleck). TGF-β, auxin, and doxycycline were reconstituted in water, while the others were reconstituted in DMSO. All drugs were stored in aliquots at -20°C.

### Time-lapse microscopy

A 24-well dish (142485; ThermoFisher) was pre-treated with 0.1% gelatin (#07903; StemCell). 2×10^4^ cells were plated 24 hr before imaging in the 24-well dish. Cells were imaged in a humidified, 37°C chamber with 5% CO2. Images were taken every 15 minutes in the cyan (excitation CWL/BW: 440/10 nm; emission CWL/BW: 460/30 nm), green (490/10 nm; 535/30 nm), red (545/10 nm; 575/25 nm), and Cy5 (610/10 nm; 670/65 nm) channels using a Nikon Ti2-E inverted microscope (Nikon). The total light exposure time was kept under 500 ms for every snapshot, and the laser intensity for each channel was kept below 5%. To quantify p21 levels, the exposure time and intensity of the cyan channel were consistently set to 200 ms and 5%, respectively, throughout all experiments.

### Image analysis

All image analyses utilized the Histone H1.0-mMaroon1 signal as a nuclear mask. We employed the automated segmentation software, Cellpose 2.0^44^, to segment individual cells within an image. For tracking single cells over time, we applied a mathematical framework known as the Linear Assignment Problem (LAP)^45^. CDK2 activity and CDK4/6 activity was calculated as the ratio between cytoplasmic and nuclear mean signals. The nuclear signal was measured within the representative region masked by H1.0-mMaroon1. The cytoplasmic signal was measured as a ring with an external diameter of 2 pixels from the nuclear boundary. CDK2 activity was used to distinguish cells in either G0/G1 arrest or proliferative state. G0/G1 arrest was defined when the mean CDK2 activity stayed below 0.8 for more than 10 hr^25^. For p21 level quantification, the mean reporter intensity within the nuclear region was measured. The 53BP1 levels were quantified by segmenting foci using watershed algorithm and counting the number of foci per nucleus.

### RNA sequencing and data analysis

Before library construction, a pellet of ∼100,000 cells were snap-frozen and stored at −80 °C. Total RNA was extracted from each sample using RNAiso Plus kit (9109; Takara). The cDNA was used for library construction using VAHTS Universal V8 RNA-seq Library Prep Kit for Illumina (NR605-01; Vazyme). The libraries were sequenced on Illumina NextSeq sequencer with PE150 chemistry. Every condition had two biological replicates. Overall sequence quality was examined using FastQC v0.11.2. Adaptor sequences were trimmed using Trimmomatic v0.32^46^. Reads were mapped to the human genome (GRCh38) using HISAT2 v2.0.3-beta^47^. Mapped fragments were counted using featureCounts^48^. Differential expression analysis was carried out using DESeq2 v1.10.1^49^. Genes with an adjusted p-value less than 0.05 and log_2_ fold change more than 1 were considered to be differentially expressed. KEGG enrichment and heatmap analysis was carried out using ClusterGVis^50^.

### Western blotting

Cells were collected and then directly lysed by addition of RIPA buffer (FD009; FUDE). Whole cell lysates were loaded onto 12% SDS-PAGE (FD346; FUDE) followed by transfer to PVDF membranes (IPVH00010; Millipore). After protein transfer, membranes were incubated in 5% milk in TBST at room temperature (RT) with rocking for at least 2 h. Primary antibody diluted in TBST was added and membranes were incubated overnight at 4°C with rocking. Membranes were washed for 5 min in TBST four times and anti-mouse (FDM007; FUDE) or anti-rabbit HRP-conjugated secondary antibodies (FDR007; FUDE) were diluted 1:10000 in TBST and incubated with membranes at room temperature with rocking for 1 hr. Membranes were washed for 5 min in TBST four times and visualized using ECL Western Blotting Substrate (E412-01; Vazyme Biotech). Blots were visualized using a gel imaging system (Clinx ChemiScope 3000 Series) and the image was analyzed by Gel-Pro analyzer software (v4.0.0.001). Antibodies used for Western blotting include p21 (CST; #2947S), GAPDH (A19056; Abclonal), mCherry (26765; Proteintech), phophos-AMPK (#2535S; CST), AMPK (66536; Proteintech), CDK6 (14052, Proteintech), CDK4 (66950; Proteintech), p53 (A195851; Abclonal), E-cadherin (20874; Proteintech), N-cadherin (22018; Proteintech), Vimentin (A19607; Abclonal), phophos-CDK2 (AP0325; Abclonal), CDK2 (A16885; Abclonal), phospho-Rb (Ser807/811) (#2524S; CST), cyclin D1 (A19038; Abclonal), cyclin B1 (A22435; Abclonal), cyclin E1 (11554; Proteintech), ERK1/2 (A4782; Abclonal) and phospho-ERK1 (T202/Y204)/ERK2 (T185/Y187) (MAB1018; R & D).

### Co-immunoprecipitation

Immunoprecipitations of p21, CDK2, and CDK6 from HL-7704 cells were performed using antibodies specific to p21 (CST; #2947S), CDK2 (A16885; Abclonal), or CDK6 (14052; Proteintech). The antibodies were captured using protein A/G magnetic beads (B23202; Selleck) according to the manufacturer’s instructions. Briefly, cells were collected and lysed directly in RIPA buffer (FD011; FUDE) on ice for 1 hour. The lysates were centrifuged at 4□°C for 10 minutes to remove cellular debris. A total of 500 μg of the cleared protein lysate was incubated with 5 μg of either rabbit IgG or the indicated antibodies at 4□°C overnight with rotation. The lysate was then incubated with 25 μL of protein A/G magnetic beads for more than 2 hr at room temperature, also with rotation. After incubation, the beads were washed with lysis buffer, and the bound proteins were eluted. The eluted samples were separated by SDS-PAGE, transferred to PVDF membranes, and probed with antibodies against p21, CDK2, CDK6, and CDK4. The antibodies used for Western blotting included p21 (CST; #2947S), GAPDH (A19056; Abclonal), CDK6 (14052; Proteintech), CDK4 (66950; Proteintech), and CDK2 (A16885; Abclonal).

### Flow cytometry

Cells were fixed with 75% ice-cold alcohol at 4°C over-night and washed three times with ice-cold PBS. Then, cells were incubated with 50 mg/ml RNase A and 40 mg/ml propidium iodide (PI) (BB-4104; Bestbio) for 30 min at room temperature. Cell cycle analysis was done by Beckman Coulter DxFLEX Cell Analyser, and the further analysis and visualization were performed in Flowjo v10.8.1 software.

### Immunostaining

Cells were fixed in an equal volume of media to 4% polyformaldehyde in PBS for 30min at 4°C. Cells were permeabilized in PBS/0.1% TritonX-100 for 15min at RT. Cells were incubated in primary antibody diluted in 5% BSA overnight at 4 °C. Cells were washed three times in PBS and then incubated with ABflo® conjugated secondary antibody (AS073, AS057;Abclonal) diluted in 5% BSA for 1 h at RT. Cells were washed three times in PBS and incubated in 1 mg/ml DAPI (P0131; Beyotime) diluted in PBS for 15 min at RT. Cells were imaged on the Nikon C2plus confocal microscope. Primary antibodies used for immunostaining in this study are: flag (AE005; Abclonal), phophos-AMPK (#2535S; CST), p27 (A0290; Abclonal).

