## Supplementary information for "Extended role of p21 during the exit from p21-induced G0/G1 arrest"

**This PDF file includes:**

Appendix Supplementary Figures 1-9

Legends for Supplementary Videos

Figure S1

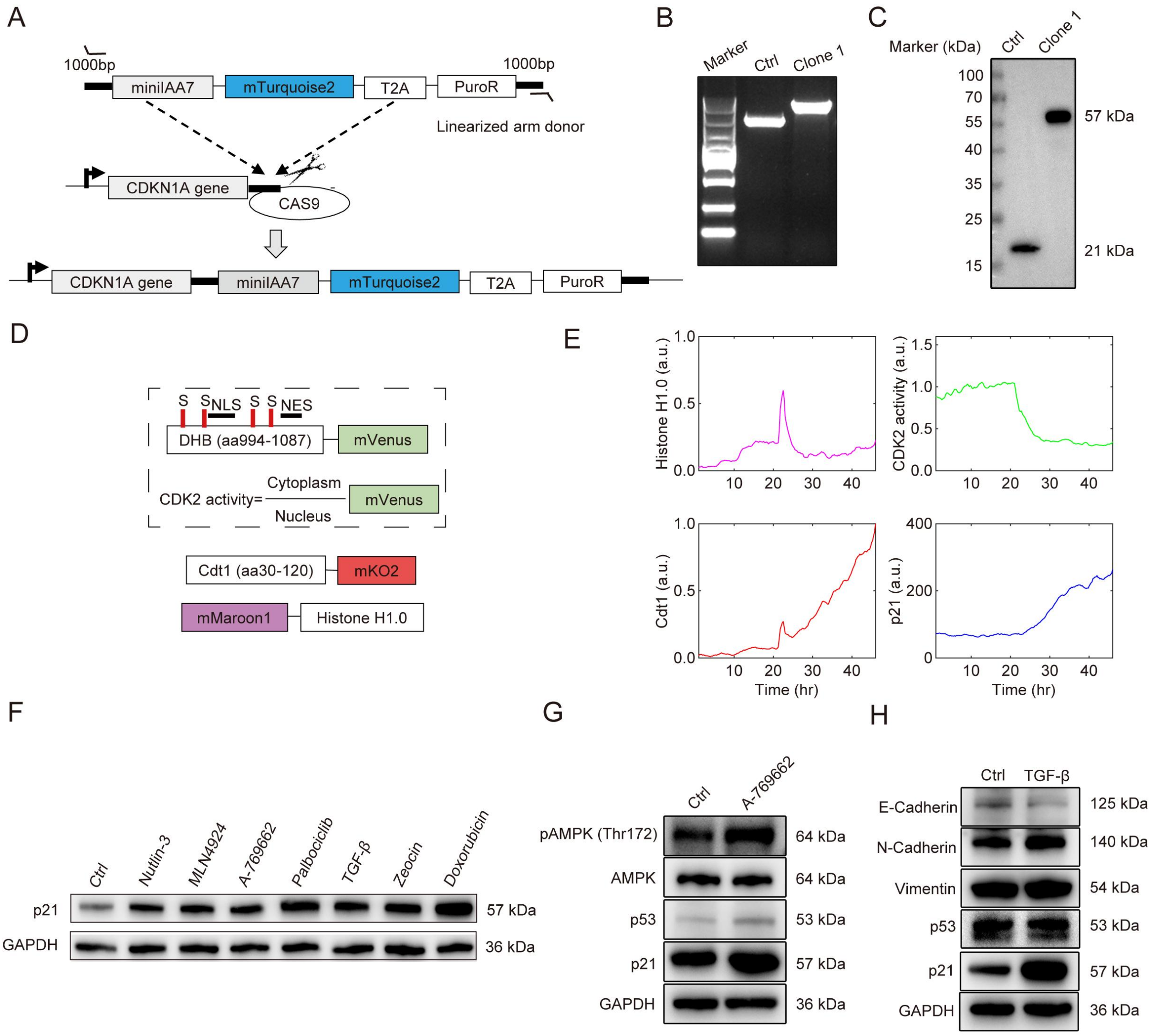


**Appendix Figure S1. System for tracking and depletion of p21 in p21-induced G0/G1 arrest.**

(A) Schematics depicting the construction of an endogenous p21-miniIAA7-mTurquoise2 knock-in cell line. Primers were designed at ~1000 bps upstream and ~1000 bps downstream of the CRISPR cutting site (black triangles). The length of the C-terminus inserted fragments (miniIAA7-mTurquoises-T2A-PuroR) was ~4000 bps, and the final p21 knock-in product expected to be ~5000 bps.

(B, C) PCR (B) and western blot (C) images showing the DNA and protein expression of homozygous p21-miniIAA7-mTurquoise2 knock-in clone (clone1) and wild type control (HL-7702 cells).

(D) Schematics showing CDK2 biosensor and Fucci reporter. CDK2-mediated phosphorylation of the serine residues in mVenus tagged DHB results in the sensor translocation from the nucleus to the cytoplasm. CDK2 activity is quantified as the ratio between the cytoplasmic and the nuclear fluorescence intensity. mKO2-tagged Cdt1 marked G1/S phase, while mMaroon1-tagged Histone1.0 serves as a nuclear marker for the nuclear segmentation.

(E) Representative single cell trajectories over 48 hr upon 5 μM Palbociclib treatment. The four trajectories showed Histone H1.0 intensity (normalized to maximum), CDK2 activity (Cyt/Nuc of DHB-mVenus), Cdt1 intensity (normalized to maximum) and p21 intensity respectively.

(F) Western blot images showing p21 and GAPDH protein level in HL-7702 p21-AID cells upon 48 hr of treatment with DMSO (control), 10 μM Nutlin-3, 50 nM Doxorubicin, 100 μM A-769662, 5 μM Palbociclib, 50 μg/ml Zeocin, 10 μg/ml TGF-β, or 1 μM MLN4924.

(G) Western blot images showing total AMPK, AMPK phosphorylation at Thr172, p53 and p21 in HL-7702 p21-AID cells treated with 100 μM A-769662 for 48 hr.

(H) Western blot images showing E-cadherin, N-cadherin, Vimentin, p53 and p21 in HL-7702 p21-AID cells treated with 10 μg/ml TGF-β for 48 hr.

Figure S2

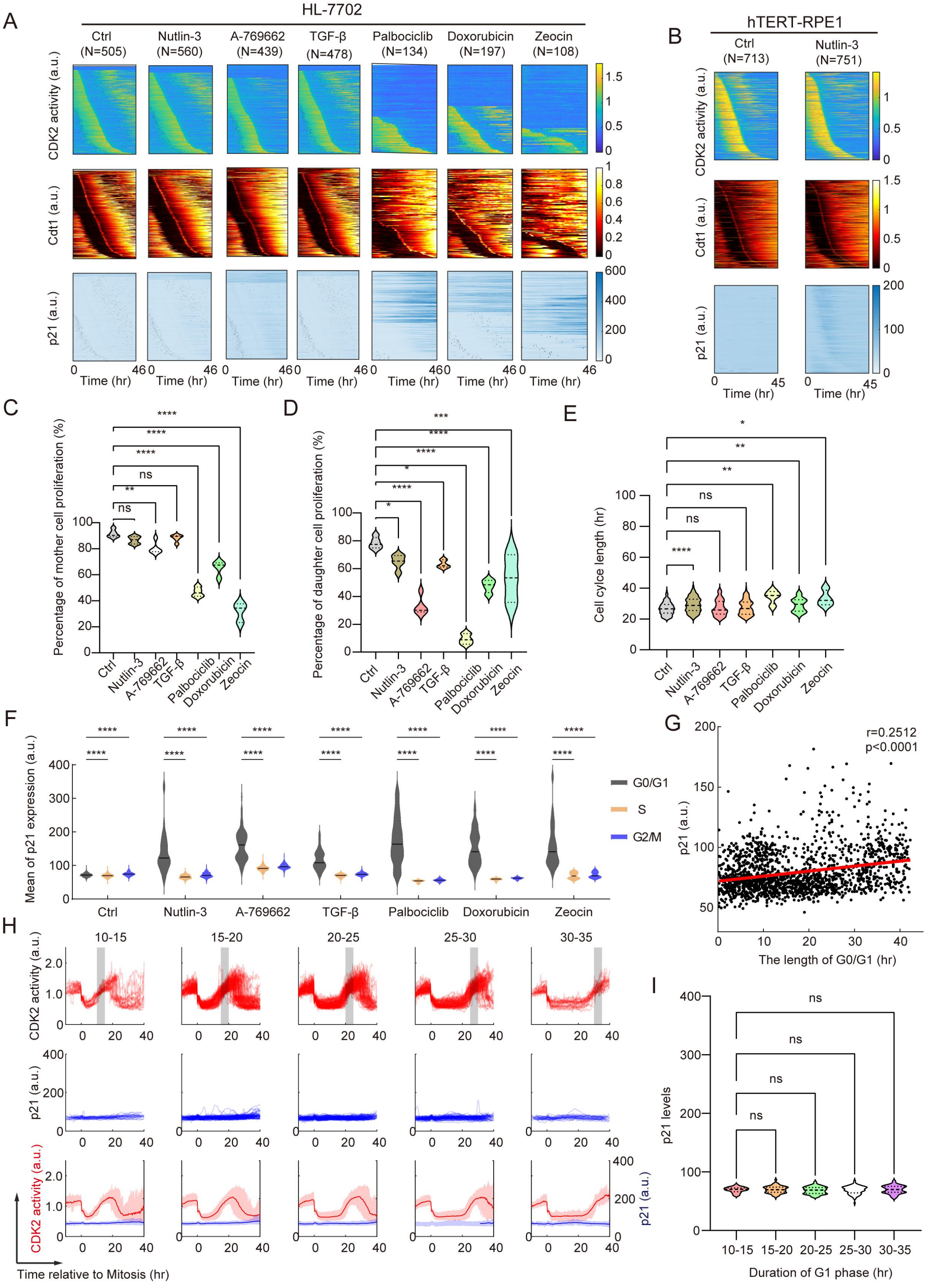


**Figure S2. The dynamic patterns of p21 expression and cell cycle under different drug-**

**induced G0/G1 arrest.**

(A) Heat maps depicting single cell dynamics of CDK2 activity, Cdt1 intensity (normalized to maximum) and p21 level in HL-7702 p21-AID cells upon DMSO (control) (*N*=505), 10 μM Nutlin-3 (*N*=560), 100 μM A-769662 (*N*=439), 10 μg/ml TGF-β (*N*=478), 5 μM Palbociclib (*N*=134), 50 nM Doxorubicin (*N*=197) or 50 μg/ml Zeocin (*N*=108) treatment. Cells were aligned based on the first time of mitotic cell division. Black dots in the p21 panel represents the timing of G1/S transition for every individual cell.

(B) Heatmaps depicting the single cell dynamic of CDK2 activity, Cdt1 intensity (normalized to maximum) and p21 in hTERT-RPE1 p21-AID cells upon DMSO (control) (*N*=713) and 10 μM Nutlin-3 (*N*=751) treatments for 45 hr. Cells were aligned based on the first time of mitotic cell division.

(C, D) Violin plot showing the percentage of proliferation in mother (C) or daughter (D) cells upon DMSO (Ctrl), Nutrlin-3, A-769662, TGF-β, Palbociclib, Doxorubicin or Zeocin treatments. One-way ANOVA was performed and "ns", *, **, *** and **** represent p-values >0.05, < 0.05, < $10^{-2}$, < $10^{-3}$ and < $10^{-4}$ respectively.

(E) Violin plot showing the length of cell cycle upon DMSO (Ctrl), Nutrlin-3, A-769662, TGF-β, Palbociclib, Doxorubicin or Zeocin treatments. One-way ANOVA was performed and "ns", *, **, *** and **** represent p-values >0.05, < 0.05, < $10^{-2}$, < $10^{-3}$ and < $10^{-4}$ respectively.

(F) Violin plots illustrating the average p21 levels at different cell cycle phases under indicated treatment conditions. **** represent p-values of Mixed-effects analysis < $10^{-4}$.

(G) Scatter plot showing the correlation between length of G0/G1 and average p21 levels in the 5 hrs' time window before p21 depletion. Pearson correlation: r = 0.2512; p-value <0.0001.

(H) Single-cell trajectories of CDK2 activity and p21 level for 80 hr during live cell imaging of HL-7702 p21-AID cells collected from all treatments. Single cells were aligned to first mitosis and categorized based on the duration of daughter cells' G1 phase. Gray shaded area represents the window of G1 phase. Bottom data are represented as mean (solid lines) ± 95% confidence intervals (shaded area).

(I) Violin plot showing the duration of G1/G0 phase and p21 levels. One-way ANOVA was performed and "ns", *, **, *** and **** represent p-values >0.05, < 0.05, < $10^{-2}$, < $10^{-3}$ and < $10^{-4}$ respectively.

### Figure S3

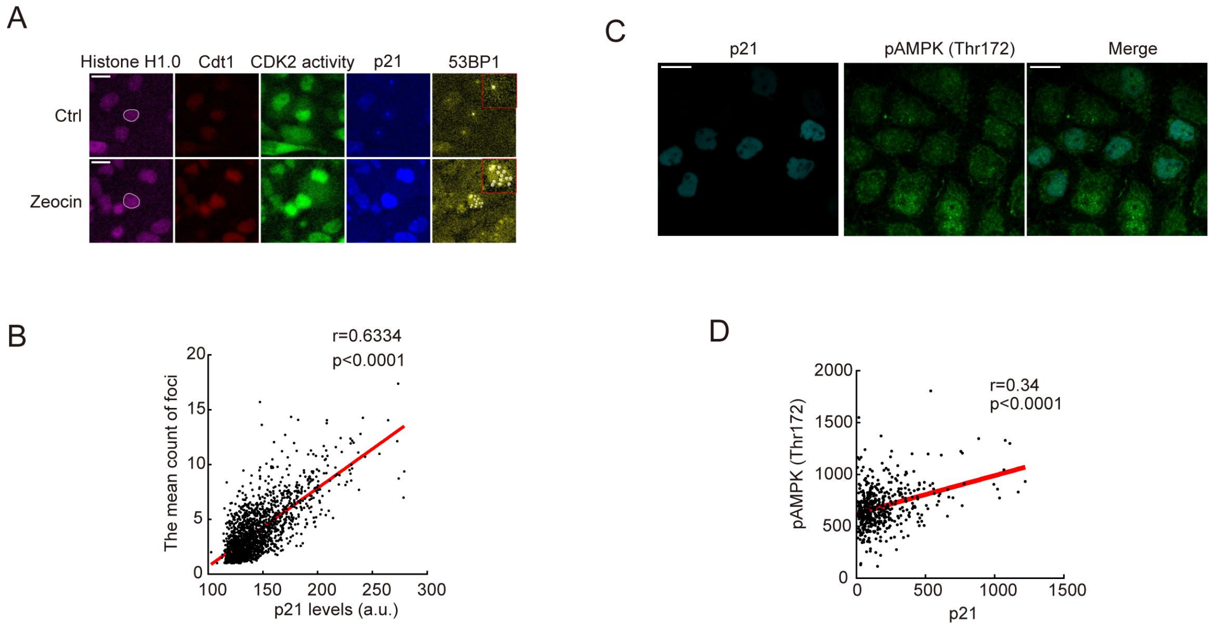


**Figure S3. The p21 level positively correlated with external stress level.**

(A) Fluorescent images showing 53BP1 reporter (truncated 53BP1 (1220-1771)) upon 1 μg/ml Zeocin treatment for 48 hr. 53BP1 reporter accumulates and forms foci representing the site of endogenous DNA damage. Scale bar, 20 μm. White circles illustrate the nuclear region of a single cell and white dots represent 53BP1 foci within the cells.

(B) Scatter plot showing the correlation between the mean 53BP1 foci count and average p21 levels measured in the last 5-hrs' time window during 48 hr Palbociclib treatment. Pearson correlation: $r = 0.6334$; p-value $< 10^{-3}$.

(C) Fluorescent images showing phosphor-AMPK (Thr172) staining in HL-7702 p21-AID cells treated with A-769662 for 48 hr. Scale bar, 20 μm.

(D) Scatter plot showing the correlation between phosphor-AMPK (Thr172) intensities measured from p21-positive cells and the corresponding p21 level. Pearson correlation $r = 0.34$; p-value $< 10^{-3}$.

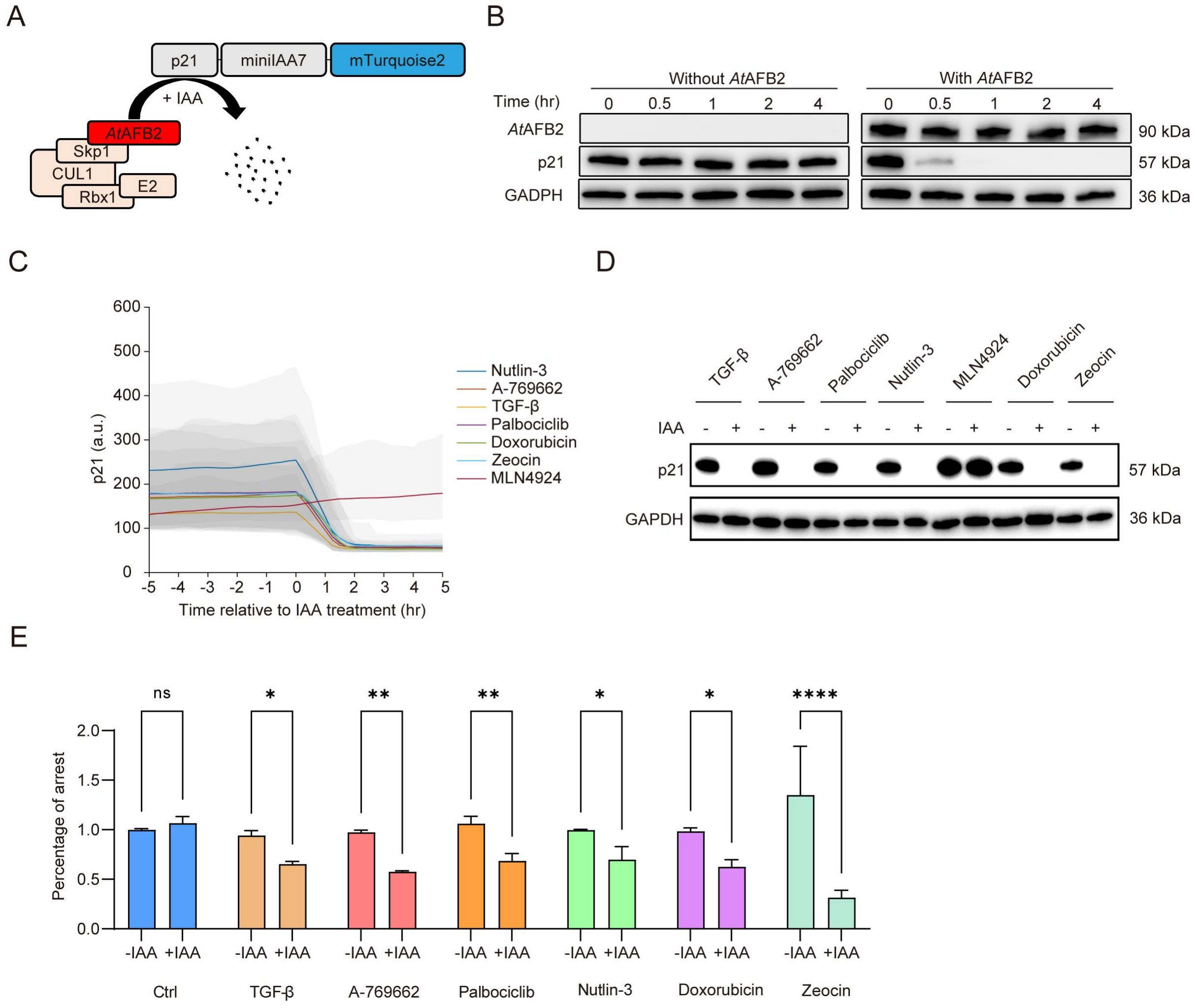


**Figure S4. p21 level and cell cycle status upon IAA treatment to deplete p21 expression.**
(A) Schematics illustrating the mechanism of p21 auxin-inducible degradation system. Under auxin treatment, the E3 ligase enzyme *At*AFB2 recognizes and ubiquitinates the miniIAA7 tag on p21, leading to p21 degradation.

(B) Western blot images showing p21, *At*AFB2 and GAPDH expression level in HL-7702 p21-AID cells with or without *At*AFB2-mCherry overexpression upon 10 μM Nutlin-3 treatment for 24 hr. Subsequently, 500 μM IAA was added to induced p21 depletion, and the cells were collected at the indicated time points.

(C) Population average trajectories showing p21 levels following the treatment of Nutlin-3, Doxorubicin, A-769662, Palbociclib, Zeocin, TGF-β, and MLN4924 with IAA addition. The conditions are aligned at the time point of IAA addition and are presented as mean (solid lines) with 95% confidence intervals (shaded area).

(D) Western blot image detecting p21 and GAPDH expression in HL-7702 p21-AID cells treated with Nutlin-3, Doxorubicin, A-769662, Palbociclib, Zeocin, TGF-β and MLN4924 for 48 hr with or without IAA addition.

(E) Bar chart showing cell cycle phases analyzed by flow cytometry upon treatment with different G0/G1 arrest inducing drugs as indicated in Figure S2A for 48 hr without or with IAA addition. IAA was added at 24 hr after drugs treatment. One-way ANOVA was performed, and "ns", *, ** and **** represent p-values >0.05, <0.05, $< 10^{-2}$ and $< 10^{-4}$. Error bars represent SEMs from three replicate experiments.

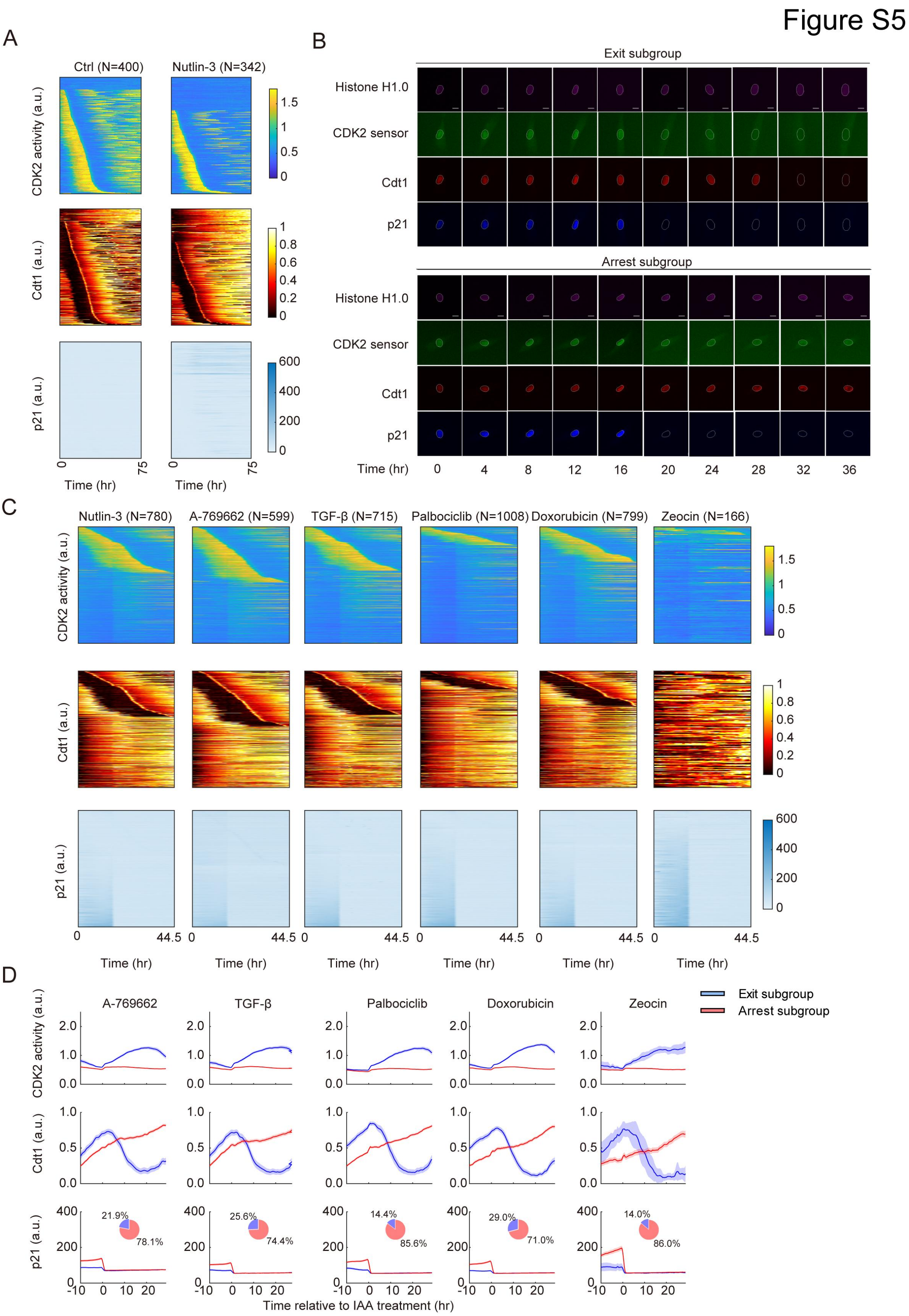


**Figure S5. The dynamics of CDK2 activity, Cdt1 and p21 under the treatment of different G0/G1 arrest inducing drugs.**

(A) Time-lapse imaging was performed to track single HL-7702 p21-AID cells treated with DMSO and Nutlin-3 for 75 hr, respectively.

(B) Single cell fluorescent live imaging showing CDK2 activity, Cdt1 and p21 expression in hTERT-RPE1 p21-AID cells following the treatment of 10 μM Nutlin-3 for 24 hr, and then cells were subjected to time-lapse imaging for 12 hr before the addition of 500 μM IAA and Nutlin-3 removal. Cells were divided into "exit" and "arrest" subgroups, based on CDK2 trajectories after IAA addition. The representative single cell fluorescent live images of CDK2 activity, Cdt1 and p21 expression. Scale bar, 10 μm.

(C) Heat maps depicting the single-cell dynamic of CDK2 activity, Cdt1 intensity (normalized to maximum) and p21 level in HL-7702 p21-AID cells under Nutrlin-3 (*N*=780), A-769662 (*N*=599), TGF-β (*N*=718), Palbociclib (*N*=1008), Doxorubicin (*N*=799) or Zeocin (*N*=166) treatments. Upon 20 hr pre-treatment with respective drugs, cells subsequently subjected to time-lapse imaging for 12 hr before the addition of 500 μM IAA and drugs removal. Cells were aligned based on the first time of mitotic cell division.

(D) Average trajectories of CDK2 activity and p21 intensity in HL-7702 p21-AID cells. Cells were treated with different drugs, including A-769662 (*N* =599), TGF-β (*N*=718), Palbociclib (*N*=1008), Doxorubicin (*N*=799), and Zeocin (*N* =166) for 24 hr, and then subjected to time-lapse imaging for 12 hr before the addition of 500 μM IAA and drugs removal. Cells were divided into two subgroups, "exit" (blue) and "arrest" (red), based on CDK2 trajectories after IAA addition. Data are presented as mean (solid lines) ± 95% confidence intervals (shaded area). Pie charts illustrate the percentage of cells in different subgroups under the indicated conditions.

Figure S6

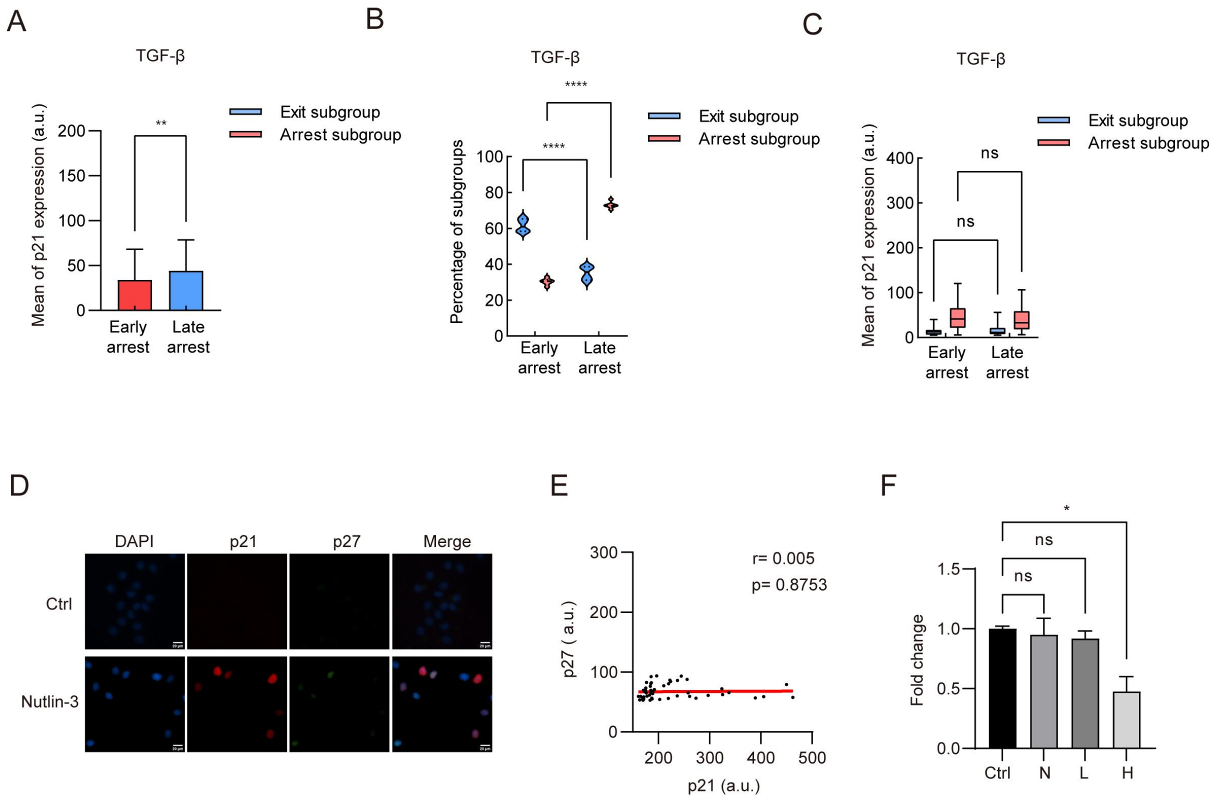


**Figure S6. p21 levels in different subgroups of cells and p27 expression in G0/G1 arrested cells.**

(A) Bar plots showing average p21 levels at early and late time points in TGF-β-induced G0/G1-arrested cells, before p21 depletion and TGF-β removal. Student's t-test was performed and ** represents p-values < $10^{-2}$. The error bars represent standard deviation (SD) for the result obtained from at least three independent live tracking experiments.

(B) Violin plots showing the percentage of the two subgroups at early and late time points following p21 depletion and TGF-β removal in cells arrested at G0/G1 by 10 μg/mL TGF-β treatment. Mixed-effects analysis was performed and **** represents p-values < $10^{-4}$.

(C) Box plots illustrating the p21 levels of the two subgroups at early and late time points following p21 depletion and TGF-β removal in cells arrested at G0/G1 by 10 μg/mL TGF-β treatment. Two-way ANOVA was performed and "ns" represents p-values >0.05. The error bars represent standard deviation (SD) for the result obtained from three independent live tracking experiments.

(D) Fluorescent images of HL-7702 p21-AID cells stained for p21 (red), p27 (green) and DAPI (blue) after treatment with or without Nutlin-3 for 48 hr. Scale bar: 20 μm.

(E) Scatter plot quantifying the correlation between p21 and p27 intensities in (D). Pearson correlation: $r = 0.005$, $p = 0.8753$.

(F) Bar plot depicting the fold change in p27 gene expression among control (Ctrl), p21 negative (N), p21 low (L), and p21 high (H) groups. Each group has at least two biological replicates. Student's t-test was performed and "ns" and * represent p-values $>0.05$ and $< 0.05$, respectively.

### Figure S7

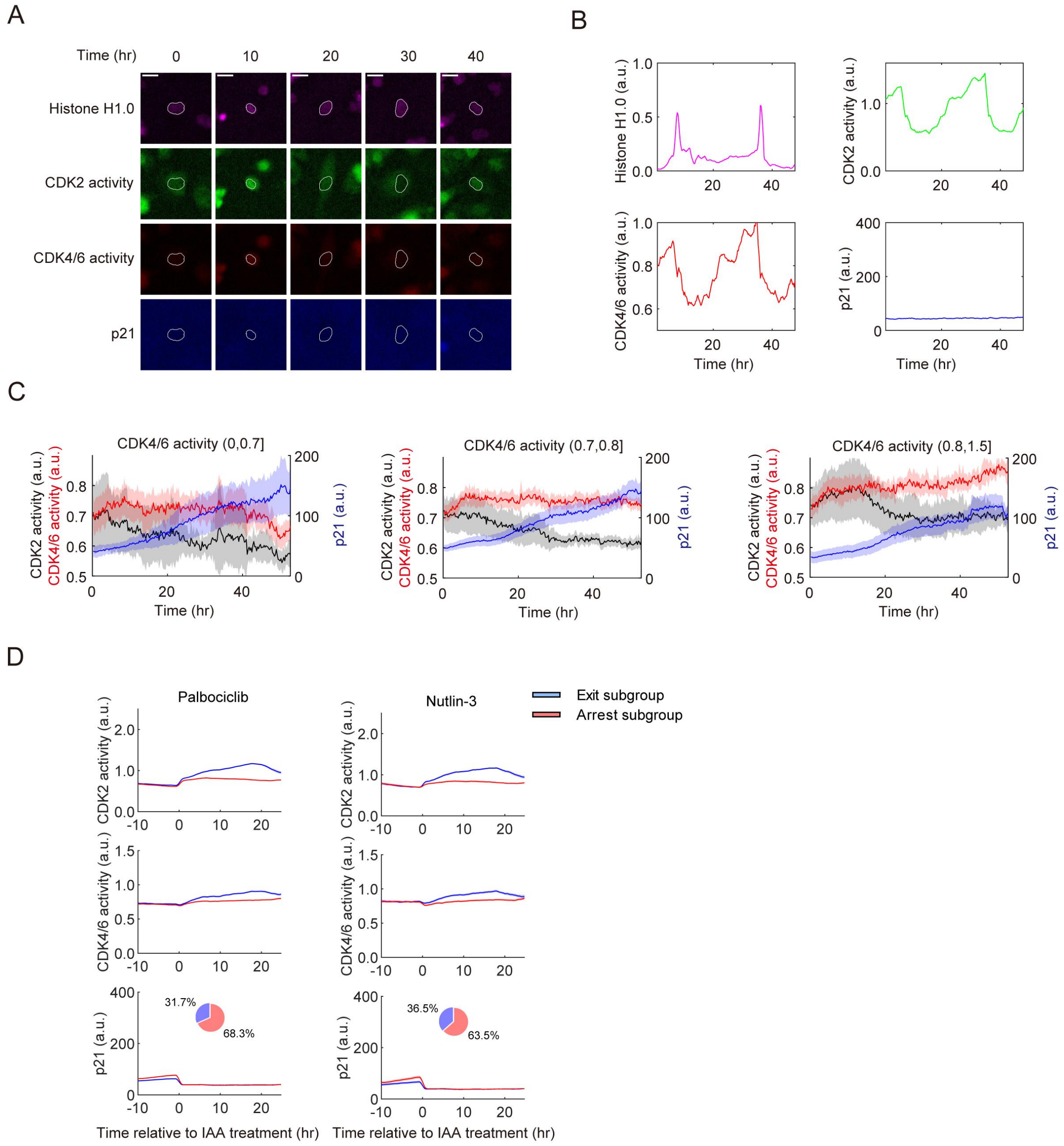


**Figure S7. The live imaging system to track the CDK4/6 sensor.**

(A) Representative single-cell fluorescent images showing CDK2, CDK4/6 and Histone H1.0 and p21 in unperturbed condition for in HL-7702 p21-AID cells stably integrated with CDK2 and CDK4/6 sensor and Histone H1.0. Scale bar, 20 μm.

(B) Representative single cell trajectories over 48 hr in unperturbed conditions. The four trajectories showed Histone H1.0 intensity (normalized to maximum), CDK2 activity (Cyt/Nuc of DHB-mVenus), CDK4/6 activity and p21 intensity respectively.

(C) Average trajectories of CDK2, CDK4/6 activity and p21 level during live cell imaging of HL-7702 p21-AID cells treated with 10 μM Nutlin-3 for 48 hr. Cells were grouped based on the average CDK4/6 activity level in the last 5-hrs' time window of the experiment. Trajectories were shown as mean (solid lines) ± 95% confidence intervals (shaded area).

(D) Average trajectories of CDK2 activity, CDK4/6 activity and p21 intensity in HL-7702 p21-AID cells. Cells were treated with Nutlin-3 (*N*=330) or Palbociclib (*N*=285) for 24 hr, and then subjected to time-lapse imaging for 12 hr before the addition of 500 μM IAA and drugs removal. Cells were divided into "exit" (blue) and "arrest" (red), based on CDK2 trajectories after IAA addition. Data are presented as mean (solid lines) ± 95% confidence intervals (shaded area). Pie charts illustrate the percentage of cells in different subgroups under the indicated conditions.

Figure S8

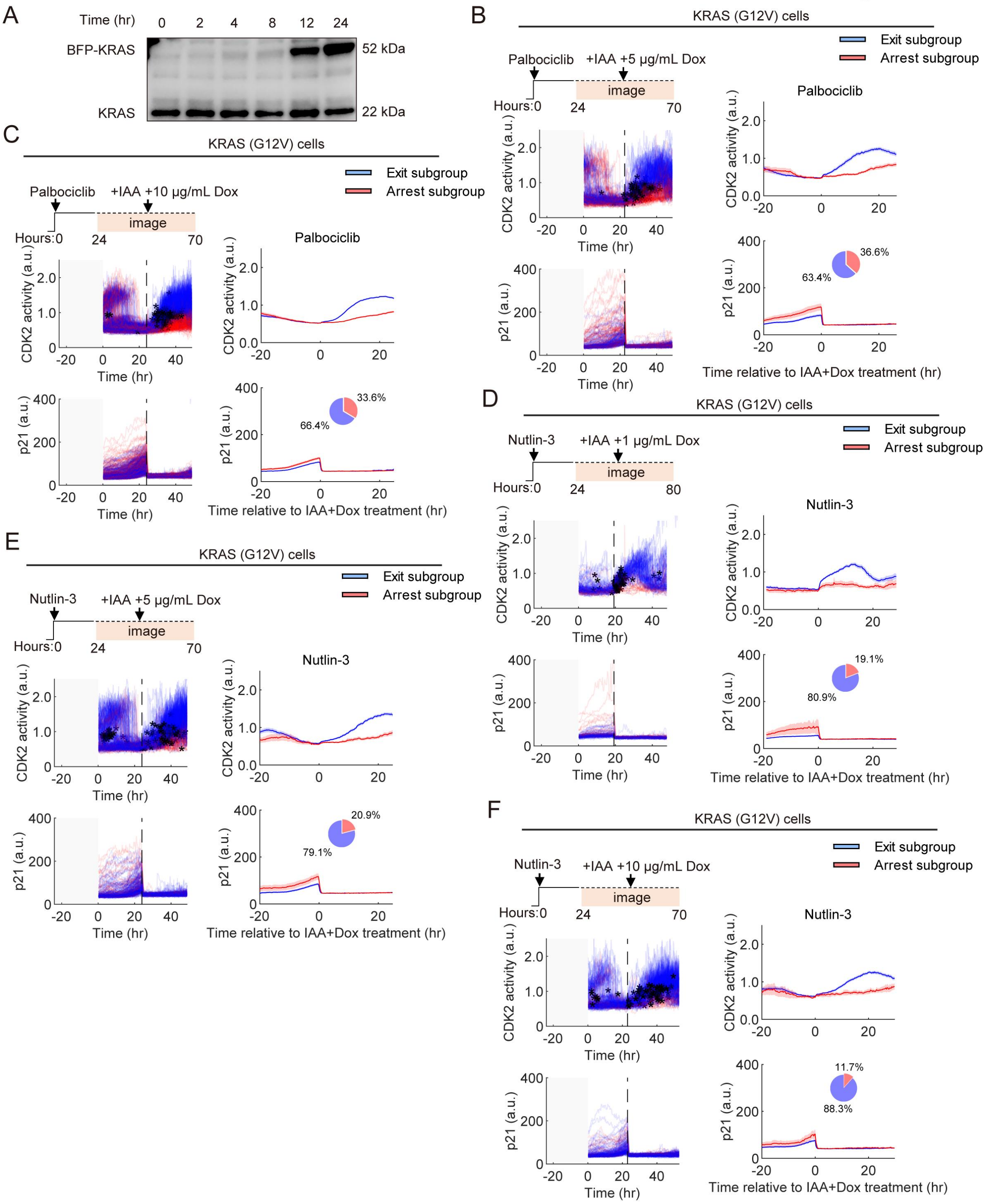


**Figure S8. RAS/ERK pathway activation rescues cell cycle re-entry defects upon arrest release without p21.**

(A) Western blot images showing the expression of endogenous KRAS and BFP-KRAS upon addition of 1μg/ml doxycycline at 0 hr in HL-7702 p21-AID cells stably expressing Tet-on KRAS (G12V).

(B, C) Single-cell trajectories (left) or average trajectories (right) of CDK2, CDK4/6 activity and p21 level during live cell imaging of HL-7702 p21-AID cells upon KRAS(G12V) overexpression after p21 depletion. Cells were pre-treated with 5 μM Palbociclib for 24 hr, and then subjected to time-lapse imaging for 12 hr before the addition of 500 μM IAA and 5 μg/ml (B, *N*= 273) or 10 μg/ml (C, *N*=884) doxycycline, together with Palbociclib removal. Vertical black arrows and dotted line indicate the time of IAA and Dox addition. Black dot denotes the time when a cell went through G1/S transition. Cells were grouped based on CDK2 activity after p21 depletion and subgroups were shown in different colors. Right panel represented mean (solid lines) ± 95% confidence intervals (shaded area). Pie plots show the percentage of the two subgroups.

(D, E, F) Single-cell trajectories (left) or average trajectories (right) of CDK2, CDK4/6 activity and p21 level during live cell imaging of HL-7702 p21-AID cells upon KRAS(G12V) overexpression after p21 depletion. Cells were pre-treated with 10 μM Nutlin-3 for 24 hr, and then subjected to time-lapse imaging for 12 hr before the addition of 500 μM IAA and, 1 μg/ml (D, *N*= 131), 5 μg/ml (E, *N*= 223) or 10 μg/ml (F, *N*= 330) doxycycline, together with Nutlin-3 removal. Vertical black arrows and dotted line indicate the time of IAA and Dox addition. Black dot denotes the time when a cell went through G1/S transition. Cells were grouped based on CDK2 activity after p21 depletion and subgroups were shown in different colors. Right panel represented mean (solid lines) ± 95% confidence intervals (shaded area). Pie plots show the percentage of the two subgroups.

### Figure S9

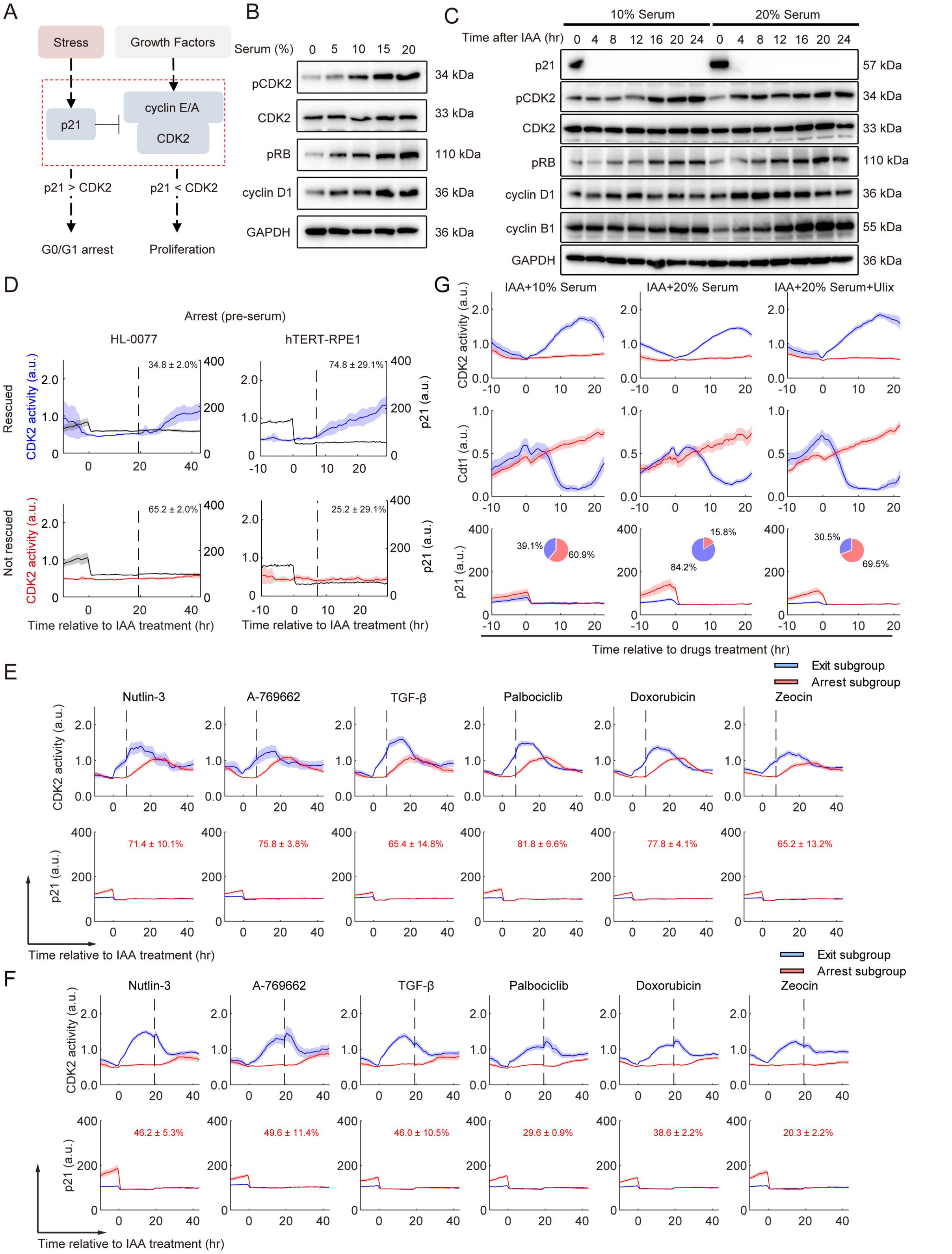


**Figure S9. High concentrations of serum rescued the cell cycle re-entry defects observed upon arrest release without p21.**

(A) Schematics depicting G0/G1 arrest-proliferation cell fate decision network under stress condition. Upon stress, p21 is transcriptionally activated, leading to inhibition of cyclin E/A-CDK2 complexes. If cyclin E/A-CDK2 complexes overcome p21 inhibition, cells proceed into proliferation. Conversely, if cyclin E/A-CDK2 complexes are inhibited by p21, cells enter a G0/G1 arrest state.

(B) Western blot images showing total CDK2, CDK2 phosphorylation at Thr160, RB phosphorylation at Ser807/811, cyclin D1 and GAPDH levels in HL-7702 p21-AID cells cultured for 24 hr in media containing 0, 5, 10 ,15 and 20% serum, respectively.

(C) Western blot images showing the level of p21, total CDK2, CDK2 phosphorylation at Thr160, RB phosphorylation at Ser807/811, cyclin D1, cyclin B1 and GAPDH levels in HL-7702 p21-AID cells. Cells were treated with 10 μM Nutlin-3 for 48 h, after which IAA was added to deplete p21 upon Nutlin-3 removal, in the presence of either 10% or 20% serum. Samples were taken at 0, 4, 8, 12, 16, 20, and 24 hr after IAA addition.

(D) Average trajectories of CDK2 activity and p21 intensity in HL-7702 and hTERT-RPE1 p21-AID cells. Cells were treated with 10 μM Nutlin-3 for 24 hr, and then subjected to time-lapse imaging for 12 hr before the addition of 500 μM IAA and Nutlin-3 removal. Subsequently, cells were treated with 20% serum at 19 hr after IAA addition, Cells were classified into "arrest" subgroups based on CDK2 trajectories after IAA addition and before 20% serum addition. Trajectories for cells that reentering or not entering cell cycle upon 20% serum were plotted individually, with the percentage shown as mean ± sd (n=3) on top. Data are presented as mean (solid lines) ± 95% confidence intervals (shaded area).

(E) Average trajectories of CDK2 activity and p21 level with high serum addition at 7 hr after p21 depletion in HL-7702 p21-AID cells. Cells were treated with Nutlin-3 ($N$= 336), A-769662 ($N$= 508), TGF-β ($N$= 290), Palbociclib ($N$= 520), Doxorubicin ($N$= 654) and Zeocin ($N$= 298), respectively, for 24 hr, and then subjected to time-lapse imaging for 12 hr before the addition of 500 μM IAA and drugs removal. Subsequently, cells were treated with 20% serum at 7 hr after IAA addition, Cells were identified as "exit" (blue) and "arrest" (red) subgroups based on CDK2 trajectories after IAA addition and before 20% serum addition. Percentages of cells that reenter cell cycle upon 20% serum treatment in the "arrest" (red) subgroups were shown on top as mean ± sd. Data are presented as mean (solid lines) ± 95% confidence intervals (shaded area). At least three replicate experiments were performed for each condition.

(F) Average trajectories of CDK2 activity and p21 level with high serum addition at 19 hr after p21 depletion in HL-7702 p21-AID cells. Cells were treated with Nutlin-3 ($N$= 381), A-769662 ($N$= 414), TGF-β ($N$= 397), Palbociclib ($N$= 344), Doxorubicin ($N$= 711) and Zeocin ($N$= 293), respectively, for 24 hr, and then subjected to time-lapse imaging for 12 hr before the addition of 500 μM IAA and drugs removal. Subsequently, cells were treated with 20% serum at 19 hr after IAA addition, Cells were identified as "exit" (blue) and "arrest" (red) subgroups based on CDK2 trajectories after IAA addition and before 20% serum addition. Percentages of cells that were rescued from the "arrest" (red) subgroups by 20% serum were shown on top as mean ± sd. Data are presented as mean (solid lines) ± 95% confidence intervals (shaded area). At least three replicate experiments were performed for each condition.

(G) Average trajectories showing CDK2, Cdt1 intensity (normalized to maximum) and p21 level during live cell imaging of HL-7702 p21-AID cells upon p21 depletion. Cells were pre-treated with 10 μM Nutlin-3 for 24 hr, followed by time-lapse imaging for 12 hr prior to addition of 500 μM IAA and Nutlin-3 removal under three conditions: 10% serum, 20% serum, or 20% serum +5 μM Ulix. Cells were divided into two subgroups based on CDK2 activity after p21 depletion with every subgroup shown in different colors. Data are represented as mean (solid lines) ± 95% confidence intervals (shaded area). Pie plots show the percentage of the two subgroups.

**Supplementary Video 1: live-cell imaging of the 53BP1 sensor in p21-AID HL-7706 cells under normal and Zeocin treatment conditions.**

The p21-AID HL-7706 cell line, expressing Histone H1.0-mMarron1 (purple), CDK2 sensor (green), Cdt1 (red), p21-mTurquoise2 (blue), and 53BP1 sensor (yellow), was imaged over 48 hr with or without 50 μg/ml Zeocin treatment. Images were captured every 15 minutes using a 20x objective lens.

**Supplementary Video 2: live-cell imaging of Nutlin-3 treated p21-AID HL-7706 cells with p21 depletion.**

The p21-AID HL-7706 cell line, expressing Histone H1.0-mMarron1 (purple), a CDK2 sensor (green), Cdt1 (red), and p21-mTurquoise2 (blue), was treated with 10 μM Nutlin-3 for 24 hr with IAA added at the 12 hr. Live cell imaging was performed over 24 hr with images acquired at 15-minute intervals using a 20x objective lens.
